# Measurement Reliability for Individual Differences in Multilayer Network Dynamics: Cautions and Considerations

**DOI:** 10.1101/2020.01.24.914622

**Authors:** Zhen Yang, Qawi K. Telesford, Alexandre R. Franco, Ryan Lim, Shi Gu, Ting Xu, Lei Ai, Francisco X. Castellanos, Chao-Gan Yan, Stan Colcombe, Michael P. Milham

## Abstract

Multilayer network models have been proposed as an effective means of capturing the dynamic configuration of distributed neural circuits and quantitatively describing how communities vary over time. Beyond general insights into brain function, a growing number of studies have begun to employ these methods for the study of individual differences. However, test-retest reliabilities for multilayer network measures have yet to be fully quantified or optimized, potentially limiting their utility for individual difference studies. Here, we systematically evaluated the impact of multilayer community detection algorithms, selection of network parameters, scan duration, and task condition on test-retest reliabilities of multilayer network measures (i.e., flexibility, integration, and recruitment). A key finding was that the default method used for community detection by the popular generalized Louvain algorithm can generate erroneous results. Although available, an updated algorithm addressing this issue is yet to be broadly adopted in the neuroimaging literature. Beyond the algorithm, the present work identified parameter selection as a key determinant of test-retest reliability; however, optimization of these parameters and expected reliabilities appeared to be dataset-specific. Once parameters were optimized, consistent with findings from the static functional connectivity literature, scan duration was a much stronger determinant of reliability than scan condition. When the parameters were optimized and scan duration was sufficient, both passive (i.e., resting state, Inscapes, and movie) and active (i.e., flanker) tasks were reliable, although reliability in the movie watching condition was significantly higher than in the other three tasks. The minimal data requirement for achieving reliable measures for the movie watching condition was 20 min, and 30 min for the other three tasks. Our results caution the field against the use of default parameters without optimization based on the specific datasets to be employed - a process likely to be limited for most due to the lack of test-retest samples to enable parameter optimization.

**Highlights:**

- Dynamic network reliability is highly dependent on many methodological decisions
- The default multilayer community detection algorithm generates erroneous results
- Reliability-optimized intra-/inter-layer coupling parameters are dataset-dependent
- Scan duration is a much stronger determinant of reliability than scan condition
- Movies are the most reliable condition, requiring at least 20 min of data

## 1. Introduction

Following early seminal contributions (Watts and Strogatz 1998, Barabasi and Albert 1999), network science has played a pivotal role in revealing the structure and interactions of complex systems, such as social and transportation networks. More recently, this methodological approach has been applied to neuroscience, helping to further characterize the architecture of the human brain and launch the field of network neuroscience (Bullmore and Sporns 2009, Bassett and Sporns 2017). Accordingly, various tools have been developed to understand the brain as a complex network, highlighting variations in brain organization across development (Gu et al. 2015), aging (Voss et al. 2013), and clinical populations (Bassett et al. 2018). In many studies, brain networks are constructed from anatomic or functional neuroimaging data as a single network or static representation (Rubinov and Sporns 2010, Sporns 2013). As the human brain is intrinsically organized into functionally specialized modules, a common approach for analyzing brain networks is to investigate community structure, which identifies areas in the brain that are densely connected internally (Sporns and Betzel 2016). While this construction is useful, a growing literature suggests that the brain, particularly its functional interactions, varies over time, thus necessitating the need to characterize these dynamic changes (Lurie et al. 2019).

Multilayer network models have been proposed as an effective means of capturing the temporal dependence between distributed neural circuits and quantitatively describing how communities vary over time (Mucha et al. 2010, Kivela et al. 2014). Multilayer network models can be used to optimize the partitioning of nodes into modules by maximizing a multilayer modularity quality function that compares edge weights in an observed network to expected edge weights in a null network. In this approach, two parameters are essential: the intra-layer coupling parameter, which tunes the number of communities within a layer, and the inter-layer coupling parameter, which tunes the temporal dependence of communities detected across layers. Dynamic network measures derived from multilayer modularity include but are not limited to flexibility, recruitment, and integration. Flexibility quantifies how frequently a region changes its community membership over time (Bassett et al. 2011); recruitment can be defined as the probability that a region is assigned to its relevant cognitive system, such as that determined by an *a priori* atlas (e.g., visual, sensorimotor, and limbic systems); and integration can be defined as the probability that a region is *not* assigned to its relevant cognitive system (Bassett et al. 2015).

Initial applications of this approach have provided key insights into the brain network dynamics that underlie learning (Bassett et al. 2011, Bassett et al. 2015). Recently, there has been increased enthusiasm to utilize these methods in the neuroimaging field (**Table 1**). Specifically, these measures have been used to link network dynamics to inter-individual differences in a broad range of functional domains, including motor learning (Bassett et al. 2011, Wymbs et al. 2012, Bassett et al. 2015, Telesford et al. 2016), working memory (Braun et al. 2015, Finc et al. 2020), attention (Shine et al. 2016), language (Chai et al. 2016), mood (Betzel et al. 2017), creativity (Feng et al. 2019, He et al. 2019), and reinforcement learning (Gerraty et al. 2018). Additionally, dynamic network reconfiguration has been suggested as a potential biomarker for diseases, such as schizophrenia (Braun et al. 2016, Gifford et al. 2020), temporal lobe epilepsy (He et al. 2018), and depression (Wei et al. 2017, Zheng et al. 2018, Shao et al. 2019, Han et al. 2020), and has been used to predict antidepressant treatment outcome (Tian et al. 2020).

**Table 1.** Summary of prior papers using flexibility, integration, or recruitment in the context of fMRI data.

| Authors | Task (Scan Duration) | Edge estimation | $\gamma$ | $\omega$ |
| --- | --- | --- | --- | --- |
| Al-Sharoha et al. (2019) | Rest (8.8 min) | Pearson's correlation coefficient | 1 | 1 |
| Bassett et al. (2011) | Motor learning (3.45 hrs) | Pearson's correlation coefficient, wavelet coherence | 1 | 1 |
| Bassett et al. (2013b) | Motor learning (3.45 hrs) | wavelet coherence | 1 | 1 |
| Bassett et al. (2015) | Motor learning (3.45 hrs) | wavelet coherence | 1 | 1 |
| Betzel et al. (2017) | Rest (10 min/session, 91 sessions) | wavelet coherence | 1 | 1 |
| Braun et al. (2015) | Working memory (~5 min) | wavelet coherence | 1 | 1 |
| Braun et al. (2016) | Working memory (~5 min) | wavelet coherence | 1 | 1 |
| Chai et al. (2016) | Semantic relatedness judgment task (13 min)<br>Story comprehension task (18~36 min) | Pearson's correlation coefficient | 1 | 0.5 |
| Cocuzza et al. (2020) | Rapid instructed task learning (8 min/run, 2 runs) | Pearson's correlation coefficient | 1 | 1 |
| Cole et al. (2014) | Dataset 1: Rest (10 min), Permuted rule operation cognitive task (72 min)<br>Dataset 2 (HCP): Rest (56 min), 7 Tasks* (total 60 min) | Pearson's correlation coefficient | 1 | 0-2 |
| Cooper et al. (2019) | Persuasive messaging task (30.3 min) | wavelet coherence | NR | NR |
| Feng et al. (2019) | Rest (~8 min) | Pearson's correlation coefficient | 1 | 1 |
| Finc et al., (2020) | Rest (10 min 10 sec/session), N-back (11 min 30 sec/session), 4 sessions | Pearson's correlation coefficient | 1 | 1, 0.5 |
| Gerraty et al. (2018) | Reinforcement learning (25 min) | wavelet coherence | 1.18 | 1 |
| Gifford et al. (2020) | Rest (5 min) | Pearson's correlation coefficient | 1 | 1 |
| Han et al. (2020) | Rest (8.5 min) | Pearson's correlation coefficient | 1 | 1 |
| He et al. (2018) | Rest (5 min), Verbal generation task (5 min) | wavelet coherence | 1 | 0.4 |
| He et al. (2019) | Rest (~8 min) | Pearson's correlation coefficient | 1 | 1 |
| Khambhati et al. (2018) | Rest (40 min) | multi-taper coherence | 1 | 1 |
| Lehmann et al. (2017) | Simulated rest (12 min) | Pearson's correlation coefficient | 1.25<br>1.5 | 2<br>1 |
| Li et al. (2019) | Rest (~6.7 min) | wavelet correlation | 1 | 1 |
| Lydon-Staley et al. (2019a) | Rest (6 min) | Pearson's correlation coefficient | 1 | 1 |
| Lydon-Staley et al. (2019b) | Rest (6 min) | wavelet coherence | 1 | 1 |
| Mattar et al. (2015) | Rest (10 min), Permuted rule operation cognitive task (72 min) | Pearson's correlation coefficient | 1 | 0.45 |
| Pedersen et al. (2018) | Rest (HCP: 60 min) | Pearson's correlation coefficient | 1 | 1 |
| Schlesinger et al. (2017a) | Dataset 1: Recognition memory task (25.5 min)<br>Dataset 2: Rest (6 min), attention task (20 min), memory task with lexical stimuli (22.5 min), face memory task (22.5 min) | wavelet coherence | 1 | 1 |
| Schlesinger et al. (2017b) | Word memory task (25.3 min) | wavelet coherence | 1.2<br>1.15 | 0.05<br>0.001 |
| Shanmugan et al. (2020) | N-back (15.4 min) | wavelet coherence | 1 | 1 |
| Shao et al. (2019) | Rest (6.75 min) | least absolute shrinkage and selection operator (LASSO) | 1 | 1 |
| Shine et al. (2016) | Rest (10 min/session, 84 sessions) | multiplication of temporal derivatives (MTD) | 1 | 1 |
| Telesford et al. (2016) | Recognition memory (20 min), Strategic attention task (20 min) | wavelet coherence | 1 | 1 |
| Tian et al. (2019) | Rest (7 min) | Pearson's correlation coefficient | 1 | 0.25 |
| Wei et al. (2017) | Rest (6.75 min) | conditional Granger causality | 1 | 1 |
| Wymbs et al. (2012) | Motor learning (3.45 hrs) | Inter-key interval (IKI) | 0.9 | 0.03 |
| Zheng et al. (2018) | Rest (8 min) | Pearson's correlation coefficient | 1 | 1 |

Note: \* 7 tasks from HCP are: emotional, gambling, language, motor, relational, social, and N-back tasks. NR: not reported.

Despite these encouraging developments, several questions remain open. First, it is unclear whether there are optimal parameter values for characterizing community structure dynamics, and the extent to which parameter choice may affect the reliability of findings. Second, the minimum data requirements to obtain reliable estimates for multilayer network-based measures have not been established. Previous studies vary in scan duration from 5 min to 3.45 hours (see **Table 1**). Third, how the choice of task during the scan (e.g., resting state, naturalistic viewing, or active tasks) impacts the reliability of dynamic network measurements obtained from multilayer modularity maximization has not been directly compared (Telesford et al., 2016). As dynamic brain network methods become more widespread, a systematic evaluation of the impact of these important factors on the test-retest reliability of multilayer network measures is important and timely (Nichols et al. 2017, Poldrack et al. 2017).

In this investigation, we aim to evaluate the impact of parameter selection, scan duration, and task condition on the test-retest reliabilities of dynamic measures obtained from multilayer modularity maximization (see **Table 2** for an overview). We first identified the optimal intra-layer and inter-layer coupling parameters for the particular multilayer community detection algorithm that we employ, based on test-retest reliability. With the optimized parameters, we then evaluated test-retest reliability at various scan durations (i.e., 10, 20, 30, 40, 50, and 60 minutes) to determine the minimum data requirements for sufficient reliability. Given the growing popularity of naturalistic viewing, we examined reliability while participants were either watching Inscapes (Vanderwal et al. 2015) or movie clips (e.g., “The Matrix”), as well as resting-state and a flanker task to directly quantify the modulatory effect of mental states. Importantly, given recent updates to dynamic community detection algorithms (Bazzi et al. 2016), we also evaluated the impact of algorithms on dynamic measurements and their test-retest reliability.

**Table 2.** Overview of the analysis performed in the paper

| Questions | Data | Analytical Approaches | Key Findings |
| --- | --- | --- | --- |
| Which modularity maximization method of the generalized Louvain algorithm is better suited for the analysis of multilayer networks in functional brain imaging data? | <u>HBN-SSI</u> (rest: 10 min/session, 12 sessions that were pseudorandomly split into a test and a retest dataset with 60-min each), n=10<br><u>HCP test-retest dataset</u> (rest: 15 min/run, 4 runs for test and 4 runs for retest), n=25<br><u>Simulated data</u> | <u>Impact of modularity maximization methods on dynamic measures</u><br>1. In HBN-SSI data, calculate three dynamic measures (flexibility, integration, and recruitment) using two modularity maximization methods (Modularity Maximization Method and Modularity Probability Method) across a range of $\omega$ [0.1 – 3.0] and $\gamma$ [0.95-1.3].<br>2. Compare two algorithms by computing the between-algorithm reliability on each of the three measures.<br>3. Replicate results in HCP and simulated data<br><u>Impact on test-retest reliability of dynamic measures</u><br>4. In HBN-SSI data, calculate the test-retest reliability for each measure and each algorithm.<br><u>Impact on validity of dynamic measures</u><br>5. In simulated data, compare the community changes obtained using the two methods and examine which one obtains results closer to the expected measures. | 1. Modularity maximization methods have a large impact on the value of dynamic network measures and these effects can be replicated in an independent neuroimaging and non-neuroimaging dataset (**Fig. 3, Fig. S1**).<br>2. Modularity Probability Method results in higher test-retest reliability for dynamic network measures (**Fig. S2**).<br>3. Modularity Probability Method recovers better the known underlying dynamics in simulated data (**Fig. S3, S4**). |
| What are the optimal parameters for test-retest reliability? | <u>HBN-SSI</u> (rest, Inscapes, movie, flanker, each with 10 min/session, 12 sessions that were pseudorandomly split into a test and a retest dataset with 60-min each), n=10<br><u>HCP test-retest dataset</u> (rest: 15 min/run, 4 runs for test and 4 runs for retest), n=25 | <u>Optimization of parameters for test-retest reliability</u><br>1. Calculate ICCs for three measures and four task conditions across the $\omega - \gamma$ plane using Craddock 200 functional parcellation.<br>2. Identify parameters resulting in peak ICC and with ICC>0.6.<br>3. Compare our parameter selection optimized for reliability ($\gamma=1.05$ and $\omega=2.5$) and previously recommended parameters ($\gamma=1$ and $\omega=1$).<br>4. Decompose ICC by assessing the within- and between-subject variability.<br><u>Comparison of optimized parameters between functional parcellations</u><br>5. Calculate ICCs across $\omega - \gamma$ plane for three measures during the movie condition using Schaefer 200 and Schaefer 600 parcellations.<br>6. Assessing within- and between-subject variability calculated using each parcellation.<br><u>Generalization of optimized parameters across datasets</u><br>7. Calculate ICCs across $\omega - \gamma$ plane for flexibility during the rest condition using HCP data | 1. We identified an optimal range of parameters generalizable across tasks and measures and the parameters peak most across tasks and measures is $\gamma=1.05$ and $\omega=2.5$ (**Fig. 4**).<br>2. Previously recommended parameters ($\gamma=1$ and $\omega=1$) resulted in high flexibility and low test-retest reliability (**Fig. S5**).<br>3. The variations in test-retest reliability on the $\omega - \gamma$ plane is driven by the within and between-subject variability (**Fig. S5, S6**).<br>4. The selection of $\omega - \gamma$ is not sensitive to functional parcellations, but to the resolution of parcellation (**Fig. S7, S8**).<br>5. Optimized $\omega - \gamma$ based on HBN-SSI data cannot be generalized to HCP data (**Fig. 5**). |
|  |  | 8. Identify parameters resulting in peak ICC and with ICC>0.6<br><u>Optimization of parameters for another reliability measure</u><br>9. Calculate discriminability for three measures and four task conditions across the $\omega - \gamma$ plane<br>10. Identify parameters resulting in peak discriminability and with discriminability>0.9. | 6. The selection of $\omega - \gamma$ is sensitive to the criteria of optimization (e.g., reliability vs discriminability) and parameters optimized for discriminability are less consistent across tasks and measures (**Fig. S9**). |
| How much data is necessary for reliably estimating multilayer network dynamics? | HBN-SSI (rest, Inscapes, movie, flanker, each with 10 min/session, 12 sessions that were pseudorandomly split into a test and a retest dataset with 10-, 20-, 30-, 40-, 50-, and 60-min each), n=10 | <u>Evaluation of ICC as a function of scan duration</u><br>1. Calculate ICCs for three measures and four tasks with 10, 20, 30, 40, 50, and 60 min of data. Each duration was repeated 100 times on randomized samples to stabilize ICC values.<br>2. Compute percent of ROIs with ICC>0.4, 0.5, 0.6<br>3. Visualize how the whole brain pattern of ICCs changes as a function of scan duration and summarize the change pattern based on Yeo et al., (2011)'s seven networks.<br><u>The impact of combining task conditions on ICCs</u><br>4. Compare ICCs obtained from 10 min of data from a single condition with those obtained from longer data created by combining data from a mixture of different conditions. | 1. The minimal data requirement for dynamic network analysis for the movie condition is 20 min and 30 min for the other three conditions (**Fig. 6, Fig.7**).<br>2. The regional and network-level variations in reliability as a function of scan duration are measure-dependent (**Fig. S10**).<br>3. When data for one task is insufficient, we can combine different tasks to increase scan duration, and thus improve reliability (**Fig. 8**). |
| Which task condition is most reliable? | HBN-SSI (rest, Inscapes, movie, flanker, each with 10 min/session, 12 sessions that were pseudorandomly split into a test and a retest dataset with 60-min each), n=10 | <u>Evaluation of ICC as a function of task condition</u><br>1. For each measure, run hierarchical linear mixed models to assess between-condition (i.e., rest, Inscapes, movie, and flanker) and between-session (i.e., test, retest) reliability in the same model.<br>2. For each measure, run simple linear mixed models for each task condition to assess between-session reliability. The significance of differences in reliability among task conditions were assessed using a nonparametric Friedman test.<br>3. For each measure, assess the within- and between-subject variability of each task condition.<br>4. Visualize how the whole brain pattern of ICCs changes as a function of task condition and summarize the change pattern based on Yeo et al., (2011)'s seven networks.<br><u>The impact of frequency on ICC during the flanker task</u><br>5. Compare ICCs obtained at low (0.01-0.1Hz) and high (0.1-0.3Hz) frequency during the flanker task. | 1. Both between-session and between-condition reliability were high for 60 min of data, except that the between-condition reliability of recruitment within the visual and sensorimotor areas is medium (**Fig. 9**).<br>2. The movie task is the most reliable (**Fig. 10**).<br>3. Regional and network-level variations in reliability as a function of task conditions are measure-dependent (**Fig. 11**).<br>4. The reliability of the flanker task is higher for the low frequency compared to the high frequency (**Fig. S11**). |

## 2. Material and methods

### 2.1 Datasets

Our primary analysis included 10 adults who had minimal head motion (median framewise displacement within 1.5 interquartile range and ranged 0.04∼0.08 mm) from the Healthy Brain Network-Serial Scanning Initiative (HBN-SSI: http://fcon_1000.projects.nitrc.org/indi/hbn_ssi/): ages 23-37 years (29.8±5.3), 50% males. HBN-SSI is a project specifically designed for evaluating the test-retest reliability of functional connectivity measures during different task states. A detailed description of the experimental design and data collection can be found in O’Connor et al. (2017). Specific details on the flanker task can also be found in our Supplementary Materials. Briefly, each participant had 12 scanning sessions collected using the same imaging protocol over a ∼1-2-month period. At each session, a high-resolution structural image and four fMRI scans (i.e., resting state, Inscapes, movie, and flanker; 10 min/condition) were collected. All imaging data were collected using a 1.5T Siemens Avanto MRI scanner equipped with a 32-channel head coil in a mobile trailer (Medical Coaches, Oneonta, NY). Structural scans were collected for registration using a multi-echo MPRAGE sequence (TR=2.73 sec, echo time=1.64 ms, field of view=256×256 mm^2^, voxel size=1.0×1.0 mm^3^, flip angle=7°). fMRI scans were collected using a multiband echo-planar imaging (EPI) sequence (multiband factor=3, TR=1.45 sec, echo time=40 ms, field of view=192×192 mm^2^, voxel size=2.46×2.46×2.5 mm^3^, flip angle=55°).

To test the impact of implementation choices on the multilayer community detection code, we included resting-state fMRI data from 25 healthy adults from the Human Connectome Project who have retest data available (https://www.humanconnectome.org/study/hcp-young-adult/data-releases) (Van Essen et al. 2013), as well as created a simulated multilayer network dataset (see Supplementary Methods for details on these datasets). Furthermore, the generalizability of parameters optimized on the HBN-SSI dataset was evaluated on the HCP retest dataset.

### 2.2 Imaging preprocessing

Functional images were preprocessed using the Configurable Pipeline for the Analysis of Connectomes (C-PAC 1.3: http://fcp-indi.github.io/) with the following steps: (1) realignment to the mean EPI image to correct for motion; (2) nuisance signal regression: regressed out linear and quadratic trends, signals of the five principal components derived from white matter and cerebrospinal fluid (CompCor, Behzadi et al. 2007), global signal (Yang et al. 2014), and Friston 24-parameter motion model (Friston et al. 1996); and (3) spatial normalization of functional data to Montreal Neurological Institute (MNI) space by combining boundary based registration (BBR) (Greve and Fischl 2009) and Advanced Normalization Tools (Avants et al. 2011). Because we are interested in the impact of both the event-related signals and the state evoked by the flanker task, we did not regress out task effects. See **Figure 1** for the flow chart summarizing the major steps of the current analytical framework.

**Figure 1.**
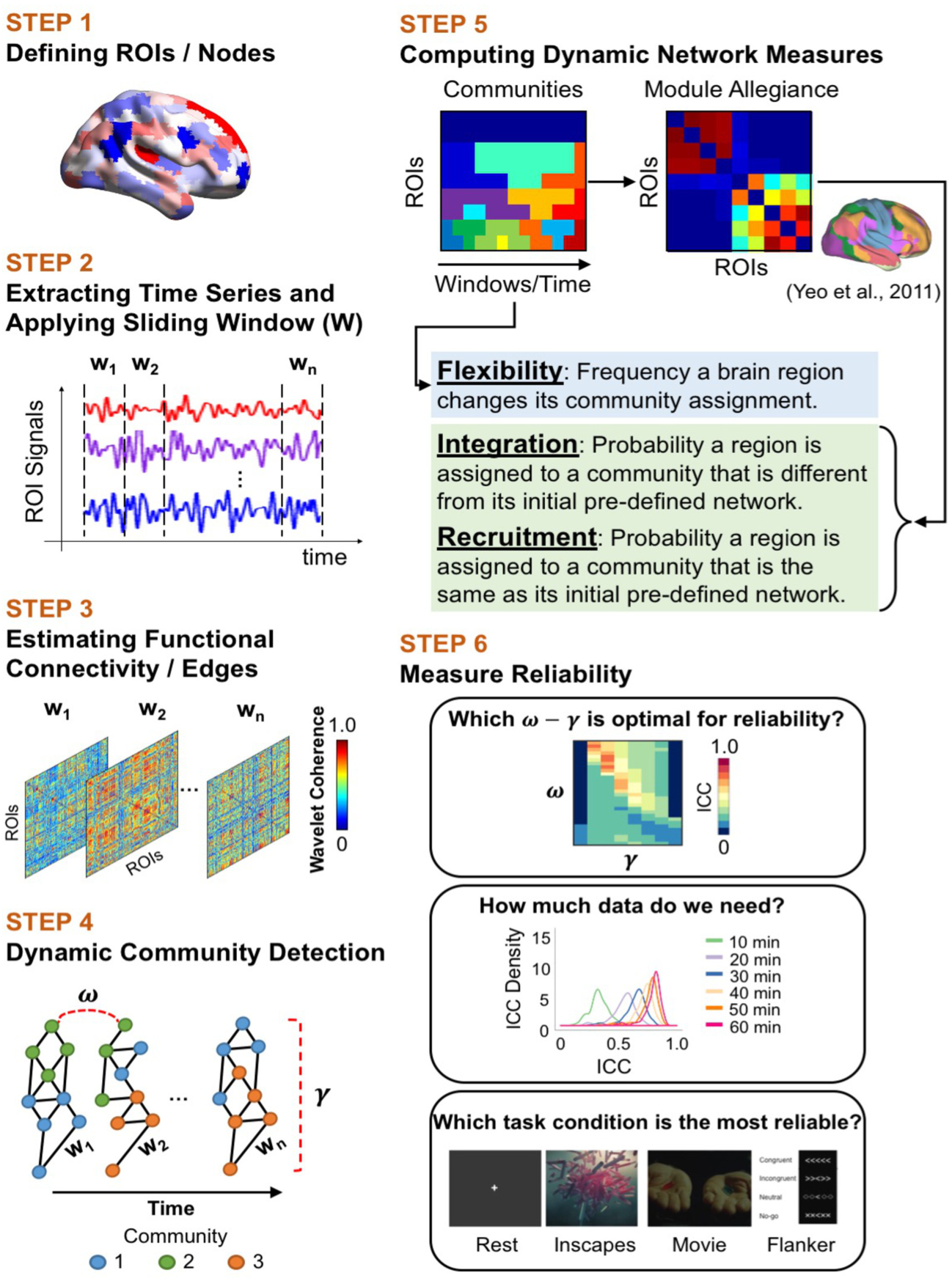
Flow chart summarizing the major steps of the current analytical framework.

### 2.3 Network construction

We defined nodes in the network using the functional parcellation from the CC200 atlas (Craddock et al. 2012) generated by a spatially constrained spectral clustering method. This functional parcellation consists of 200 ROIs covering the whole brain, each of which is homogeneous in its estimated functional connectivity. This commonly chosen atlas was previously used for studying static functional connectivity in the same dataset (O’Connor et al. 2017) and for evaluating the reproducibility and reliability of state-based temporal dynamic methods (Yang et al. 2014). To determine whether our results were sensitive to functional parcellations and the resolution of parcellations, we tested the robustness of our findings using the Schaefer 200 and 600 brain parcellations (Schaefer et al. 2018). After preprocessing, we extracted mean signals from each ROI and then applied a sliding window to the time series. The window length (∼100 s, 68 TRs, no overlap) was selected based on a previous multilayer network study (Telesford et al., 2016), which demonstrated that the number of communities stabilizes at a window length of ∼100 s and that the inter-region variance of flexibility peaks at a window size of 75∼120 s across different cognitive tasks. Since we are interested in comparing the test-retest reliabilities of dynamic network measures among task conditions, we selected the window length of ∼100 s to also capture low frequency fluctuations with a low cutoff at 0.01 Hz. However, we acknowledge that this window selection may not have sufficient temporal resolution to relate network dynamic changes to changing conditions in naturalistic viewing or in the flanker task.

For each window or layer, edges were estimated using wavelet coherence using the wavelet coherence toolbox (Grinsted et al. 2004) (http://grinsted.github.io/wavelet-coherence/). As the most commonly used edge estimation for multilayer network analyses (**Table 1**), wavelet coherence is robust to outliers (Achard et al. 2006) and has advantages in terms of its utility for estimating correlations between fMRI time series, which display slowly decaying positive autocorrelations or long memory (Zhang et al. 2016, Telesford et al. 2017). Specifically, magnitude-squared coherence *C*_*xy*_ between a given pair of regions (x, y) is a function of the frequency (*f*) and defined by the equation:

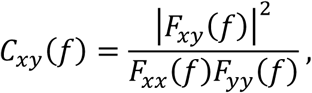

where *F*_*xy*_(*f*) is the cross-spectral density between region x and region y. The variables *F*_*xx*_(*f*) and *F*_*yy*_(*f*) are the autospectral densities of signals from region x and region y, respectively. The mean of *C*_*xy*_(*f*) over the frequency band of interest, in our case 0.01-0.10 Hz, is the edge weight between regions x and y. The range of wavelet coherence is bounded between 0 and 1. For each subject, we obtained a 200×200×6 (region×region×window) coherence matrix per task per session, which is coupled into a multilayer network by linking a node to itself in the preceding and the following windows or layers (Mucha et al. 2010, Bassett et al. 2011). Dynamic community detection was then performed for each session.

### 2.4 Dynamic community detection algorithm

Multiplex communities can be defined by modularity maximization (Mucha et al., 2010), spectral clustering (Lin et al. 2009, Michoel and Nachtergaele 2012), or other data-mining approaches (Ströele et al. 2009, Ströele et al. 2011, Ströele et al. 2012). Among these methods, the optimization of multislice modularity is the most popular approach for fMRI research, possibly due to its feasibility of representing dynamic functional networks (Bassett et al. 2011, Bassett et al. 2013a, Bassett et al. 2013b). While there are numerous methods for determining community structure, here we used a Louvain-like locally greedy algorithm (Blondel et al. 2008). We chose this algorithm because: (1) it has been shown to outperform other community detection methods (Yang et al. 2016), (2) is most commonly used in the field of network neuroscience (Blondel et al. 2008), (3) has been adapted to multilayer network models (Mucha et al. 2010, Bassett et al. 2011, Bassett et al. 2013a), and (4) has been commonly used in studies linking multilayer network measures to cognition and disorders. For detecting communities, the multilayer modularity quality function (Q) is optimized (details given in section 2.4.1) and is defined as in Mucha et al. (2010):

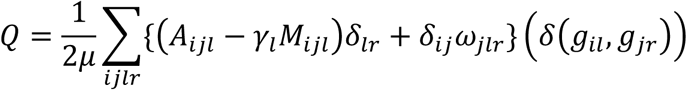

where μ is the sum of edge weights across all nodes and layers; δ_ij_ is the Kronecker’s δ-function that equals 1 when i = j and equals 0 otherwise. The element A_ijl_ gives the strength of the edge between nodes *i* and *j* in layer l, and the element M_ijl_ is the corresponding edge expected in a null model. Different choices of the null model in modularity could lead to different community structures (Sarzynska et al. 2016); based on the scope of the current paper, we adopted the widely used setting in the dynamic network reconfiguration papers: the Newman-Girvan null model which defines M_ijl_ as:

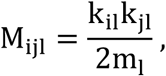

where 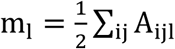 is the total edge weight in layer l. The variables k_il_ and k_jl_ are the intra-layer strengths of node *i* and node *j* in layer l, respectively. In the quality function, g_il_ represents the community assignment of node *i* in layer l, and g_jr_ represents the community assignment of node *j* in layer *r*. Finally, δ(g_il_, g_jr_) = 1 if g_il_ = g_jr_ and δ(g_il_, g_jr_) = 0 if g_il_ ≠ g_jr_.

We performed multilayer community detection using the generalized Louvain package implemented in MATLAB (Lucas et al. 2011-2019). This method treats intra-layer and inter-layer edges as unique and assigns communities to regions in all layers. This allows for the investigation of communities that are coherent over time and simultaneously across layers. Moreover, as the community labels are consistent across layers, this avoids the common problem of community matching.

#### 2.4.1 Algorithm selection

Optimization of the quality function or modularity (Q) includes two phases: community detection and community merging (**Figure 2**). In the first phase, each node starts as its own community. Starting from a randomly chosen initial node, modularity is calculated after merging this node with every other community, one by one. Then all merges that increase modularity are identified.

**Figure 2.**
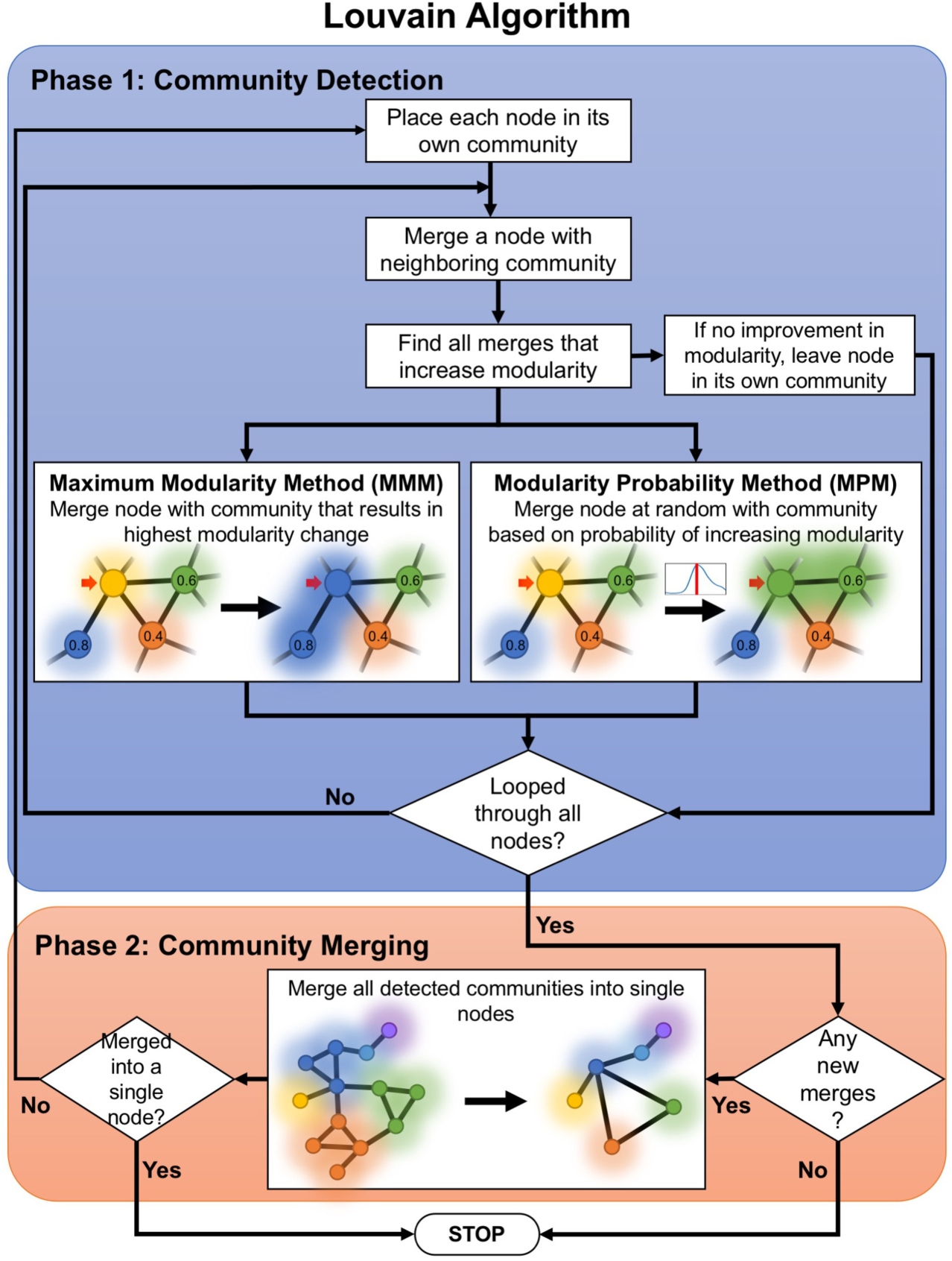
Diagram showing the modularity maximization process. The generalized Louvain algorithm is a two-phase process that finds communities in a network. In the first phase, every node is placed in its own community. At random a node is merged with a neighboring community and modularity is calculated; after iterating through all available communities, a node can be merged with a community based on different methods. In the Maximum Modularity Method (MMM), the merge that resulted in the greatest increase in modularity is chosen. In the Modularity Probability Method (MPM), a community is chosen at random based on the probability that it increases the modularity. This process continues sequentially for all nodes and/or until there are no improvements (no increase in modularity). In the second phase, detected communities are merged into single nodes and the process is repeated again. The algorithm ends if all nodes merge into a single community or if there are no improvements in modularity after iterating through nodes.

The Default algorithm (Maximum Modularity Method: MMM) and the Revised algorithm (Modularity Probability Method: MPM) differ in how the merge is selected. The MMM selects the merge that produces the highest increase in modularity. In the MPM, a probability is attached to all merges that increase modularity (the higher the proportion of modularity increases, the higher the probability). Afterward, a merge is chosen randomly, weighted by the probability distribution of modularity increases. If no improvement in modularity is found, the node is left unmerged. This process is then repeated sequentially for all other nodes. In the second phase, any multi-node community is merged and treated as a single node. Then the two phases are repeated until all communities are merged into a single community or no further improvement is possible.

The MMM was the default method implemented in the original code publicly released in 2011. The MPM was added in 2016 (Version 2.1) to address an abrupt change in the behavior of the default method when the inter-layer coupling parameter increased (see Bazzi et al. 2016 for details). This abrupt change was initially observed in financial data and has not been evaluated in neuroimaging data. Given that the default method was widely used in the fMRI literature, we evaluated the impact of the default and the improved method on the values of dynamic network measures, as well as the reliability and validity of these measures before making a selection.

#### 2.4.2 Parameter optimization

When optimizing multilayer modularity, we must choose values for the two parameters γ and ω. The parameter γ_l_ is the intra-layer coupling parameter for layer l, which defines how much weight we assign to the null network and controls the size of the communities detected within layer l. The parameter ω_jlr_ is the inter-layer coupling parameter, which defines the weight of the inter-slice edges that link node *j* to itself between layer l and layer *r*; this parameter controls the number of communities formed across layers. Here, following previous work in the neuroimaging literature (**Table 1**), these two parameters have been set as constants (γ_l_=γ and ω_jlr_=ω) across layers. The choice of these two parameters is critical for multilayer modularity optimization, as they have a large impact on the detected community structure, as well as on the dynamic measures derived from multilayer communities (Bassett et al. 2013a, Mattar et al. 2015, Chai et al. 2016). Multilayer modularity approaches were also shown to detect spurious group differences in dynamic network measures when these parameters were set inappropriately (Lehmann et al. 2017). Here, we optimized these two parameters based on test-retest reliability. Specifically, we computed intra-class correlation coefficients (ICC) for each of the three dynamic network measures across a range of γ and ω for each of the four tasks. Specifically, we considered the space spanned by the following ranges: γ=[0.95, 1.3] and ω=[0.1, 3.0]. We determined these ranges by applying the criterion that the number of modules be ≥2 and ≤100. As the space for γ is much smaller than that for ω, a smaller increment of 0.05 was used for γ and an increment of 0.1 was used for ω. After estimating the ICC at each point in this space, we identified the parameter value pair that produced the largest ICC. The γ and ω pair that produced the largest ICC most frequently across the 12 conditions (3 dynamic network measures and 4 tasks) was chosen as the optimal one for the dataset.

In addition to ICC, we also used the discriminability to quantify the reliability and optimize the parameters for multilayer network measurement. Discriminability is a non-parametric statistic to quantify the degree to which an individual’s samples are relatively similar to one another, which can be used to measure the reliability without restricting the data to be univariate and Gaussian distribution (Bridgford et al. 2019). Here, we computed the discriminability across the γ-ω plane for flexibility, integration, and recruitment separately. Considering n subjects, where each subject has s measurements, then we have N =n×s total measurements across subjects for a given dynamic measure. Then discriminability is computed in the following three steps: (1) Compute the distance (in this case the Euclidean distance) between all pairs of measurements, resulting in a N×N matrix; (2) For measurements of all subjects, compute the fraction of times that a within-subject distance is smaller than a between-subject distance, resulting in N×(s-1) numbers between 0 and 1 (Craddock et al. 2012). The discriminability of the dataset is the average of the above mentioned fractions across subjects, resulting in a single number between 0 and 1. A high discriminability indicates that within-subject measurements are more similar to one another than between-subject measurements, suggesting the measurement is more reliable. The calculation of discriminability is conducted using R package (https://github.com/ebridge2/Discriminability)

#### 2.4.3 Other considerations

When implementing the GenLouvain method, we used fully weighted, unthresholded coherence matrices to minimize the known near degeneracy of the modularity landscape (Good et al. 2010). After applying this algorithm, the 200 ROIs were assigned to communities that spanned across layers. Due to the roughness of the modularity landscape (Good et al. 2010) and the stochastic nature of the algorithm (Blondel et al. 2008), the output of community detection often varies across optimizations. Thus, rather than focus on any single optimization, we computed the dynamic measures based on 100 optimizations, following the precedent of previous work (Bassett et al. 2011, Bassett et al. 2013a, Bassett et al. 2013b, Bassett et al. 2015). Specifically, we first calculated network measures (see next section for details) for each run of the community detection algorithm, and then we averaged those measures over the 100 optimizations.

### 2.5 Calculation of dynamic network measures

For each participant, we computed the following measures to characterize the dynamics of the multilayer network based on the dynamic community structure detected in each optimization.

#### 2.5.1 Flexibility

For each brain region, the flexibility is calculated as the number of times a brain region changes its community assignment across layers, divided by the number of possible changes, which is the number of layers minus 1 (Bassett et al., 2011). This measure characterizes a region’s stability in community allegiance and can be used to differentiate brain regions into a highly stable core and a highly flexible periphery (Bassett et al. 2013b). Regions with high flexibility are thought to have a larger tendency to interact with different networks. Average flexibility across the brain is also computed to examine the global flexibility of the system.

#### 2.5.2 Module allegiance

The module allegiance matrix is the fraction of layers in which two nodes are assigned to the same community (Bassett et al. 2015). For each layer, a co-occurrence matrix (200×200) can be created based on the community assignment of each node pair. The element of the co-occurrence matrix is 1 if two nodes are assigned to the same community, and 0 otherwise. The module allegiance matrix is computed by averaging the co-occurrence matrices across layers, and the value of the matrix elements thus ranges from 0 to 1.

#### 2.5.3 Integration and recruitment

To quantify the dynamic functional interactions among sets of brain regions located within predefined functional systems (i.e., seven networks defined by Yeo et al. 2011), we computed two network measures based on the module allegiance matrix: recruitment and integration (Bassett et al., 2015). Recruitment can measure the fraction of layers in which a region is assigned to the same community as other regions from the same pre-defined system. The recruitment of region i in system S is defined as:

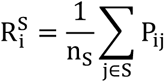

where N_S_ is the number of regions in S, and P_ij_ is the module allegiance between node i and node j. The integration of region i with respect to system S is defined as:

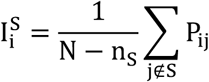

where N is the total number of brain regions. Integration 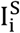 measures the fraction of layers in which region i is assigned to the same community as regions from systems other than S.

### 2.6 Assessment of reliability

Test-retest reliability and between-code reliability were assessed with the ICC estimated using the following linear mixed model:

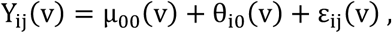

where Y_ij_(v) represents the dynamic measure (i.e., flexibility, integration, or recruitment) for a given brain region v (v=1, 2…, 200), *i* indexes participants (i=1, 2, … 10), and *j* indexes either the session for analyses of test-retest reliability or the code implementation options for analyses of between-code reliability (j=1, 2). Further, μ_00_(v) is the intercept or a fixed effect of the group average dynamic measure at region v; θ_i0_(v) is the random effect for the i-th participant at region v; and ε_ij_(v) is the error term. The total variance of a given dynamic measure can be decomposed into two parts: (1) inter-individual variance across all participants 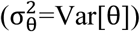, and (2) intra-individual variance for a single participant across two measurements 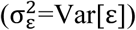. The reliability of each dynamic measure can then be calculated as:

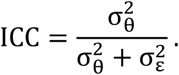

The model estimations were implemented using the linear mixed effect (lme) function from the nlme R package (http://cran.r-project.org/web/packages/nlme).

### 2.7 Determination of the minimal data requirement

To establish minimal data requirements for sufficient test-retest reliability, we compared ICC values of six scan durations: 10 min, 20 min, 30 min, 40 min, 50 min, and 60 min. Different scan durations were obtained by pseudo-randomly selecting 1, 2, 3, 4, 5, or 6 10-min sessions from 12 available sessions for each participant. Dynamic features were first computed for each of the 12 10-min sessions, and then averaged across the sessions that were selected for each scan duration. We did not compute the dynamic measures on concatenated time series data to avoid artifactually introducing community changes at the concatenation point. For each scan duration, ICC was estimated using linear mixed models. To increase the robustness of the results and to extract stable features, we repeated the analysis on 100 randomized samples for each duration. The same process was performed for each of the four tasks to determine the data necessary for each condition.

### 2.8 Determination of task dependency

To investigate how estimates of test-retest reliability might depend on task states, we first used hierarchical linear mixed models to assess between-condition and between-session reliability in the same model. Hierarchical linear mixed models separate the variations among task conditions (i.e., between-condition reliability) from variations between sessions (i.e., test-retest reliability) by estimating variance between participants, across the four task conditions (for the same participant), and between sessions within each condition (O’Connor et al. 2017). Our model took the following form:

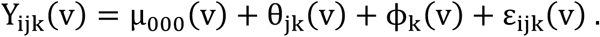

The dynamic measure for a given brain region v can be denoted as Y_ijk_(v), where *i* indexes over sessions, j indexes over conditions, and k indexes over participants. In this model, μ_000_ represents the intercept; θ_jk_ represents a random effect between sessions for the j-th condition of the k-th participant; ϕ_k_ represents a random effect for the *k*-th participant; and ε_ijk_ represents the error term. The variables θ_jk_, ϕ_k_, and ε_ijk_ are assumed to be independent and to follow a normal distribution with a zero mean. The total variances of a given dynamic measure can be decomposed into three parts: (1) variance between participants 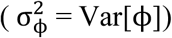; (2) variance between conditions for the same participant 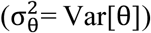; and (3) variance of the residual, indicating variance between sessions 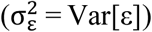. The reliability of each dynamic measure across conditions can be calculated as

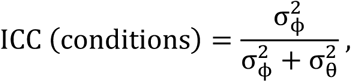

and across sessions as

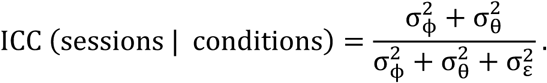

Next, we estimated the test-retest reliability for each task using the simple linear mixed models described in Section 2.7. The main effect of task condition on ICC values was tested using a nonparametric Friedman test. The Wilcoxon signed-rank test was used for *post hoc* analyses to determine which tasks differed significantly in test-retest reliability. As ICCs consistently increase with scan duration (Laumann et al. 2015, Xu et al. 2016, O’Connor et al. 2017), hierarchical and simple linear mixed models were performed using 60 min of data (the optimal scan duration in the current sample) to determine the impact of task condition.

## 3. Results

### 3.1 Impact of modularity maximization algorithm

In 2016, a comprehensive examination of multilayer networks with financial data revealed an abrupt discontinuity in values across the γ-ω landscape when the Maximum Modularity Method (MMM; the default method) was used. These findings raised concerns about the robustness of MMM (see Figure 5.4 in Bazzi et al. 2016). In our study, when we used MMM, we observed a similar discontinuity in multilayer network-based dynamic measures in two independent human brain imaging datasets (HBN-SSI and HCP), as well as in a simulated multilayer-network dataset (**Figure 3**). When the updated Modularity Probability Method (MPM) was used, we no longer observed such apparent discontinuities. To compare the dynamic measures computed using these two methods, we assessed the between-method reliability of flexibility for the two algorithms. Consistent with our intuition, we found that most of the ICC values above the discontinuity were near zero, suggesting that flexibility values obtained using different randomization methods can differ dramatically in that portion of the parameter space. In addition to flexibility, we also investigated the impact of these two methods on integration and recruitment. We found that flexibility was the most impacted, integration was impacted less, and recruitment was the least impacted (**Figure S1**). Furthermore, we found that the updated method produced measures with greater test-retest reliability than the default method (**Figure S2**), and better recovered known underlying dynamics in the simulated data - especially in the portions of parameter space above the apparent discontinuity (See **Figure S3** and **S4** for details). Thus, the updated method (MPM) was used in the present work.

**Figure 3.**
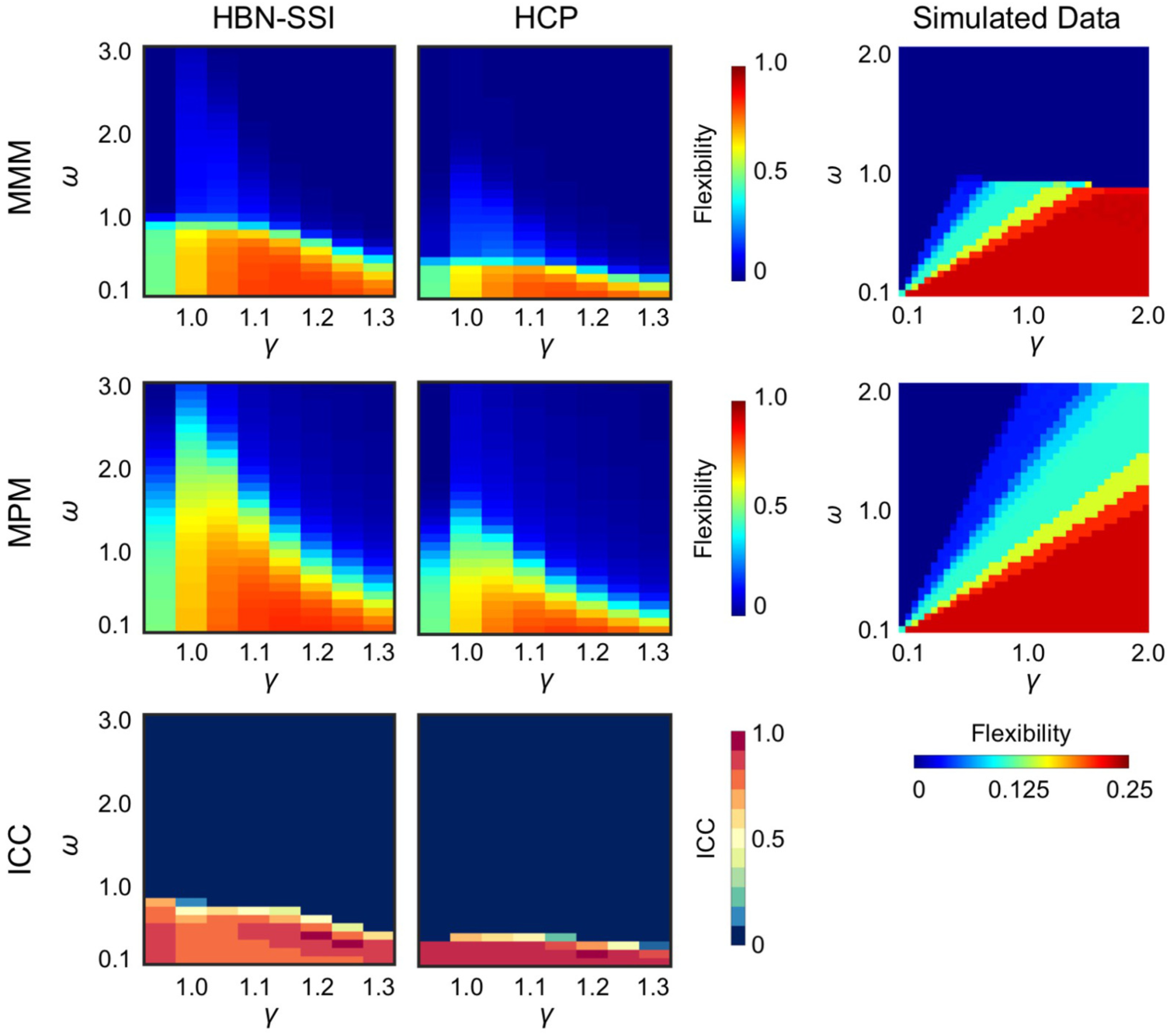
The impact of generalized Louvain method on estimated values of flexibility. When the default MMM was used, there was a dropoff in flexibility values in the 2-dimensional γ-ω parameter space (Top row). This apparent discontinuity was observed on flexibility values in two independent human brain imaging datasets, the Healthy Brain Network-Serial Scanning Initiative (HBN-SSI) and the Human Connectome Project, as well as in simulated data. The issue was mitigated by the updated MPM (Middle row). Reliability between the two randomization methods quantified using intra-class correlation coefficients (ICCs) was good below the apparent discontinuity and was near zero above the discontinuity (Bottom row). ICCs for HBN-SSI and HCP were evaluated based on 60 min of resting state data.

**Figure 4.**
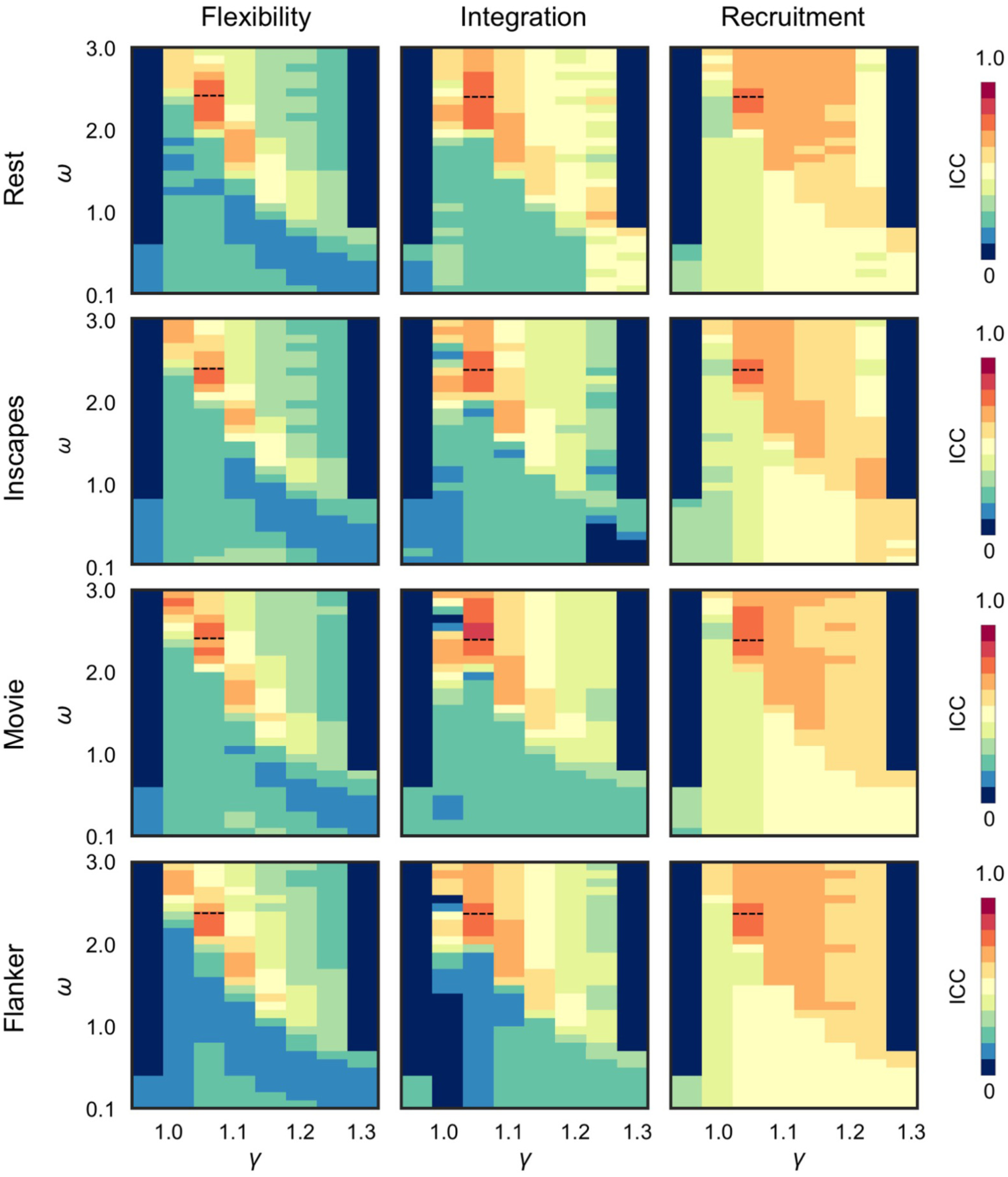
Test-retest reliability of dynamic network measures depends on the γ-ω selection for the Healthy Brain Network-Serial Scanning Initiative (HBN-SSI) dataset. In the γ-ω plane, global ICC was computed across 200 ROIs. We identified a range of parameters that produced good test-retest reliability (ICC≥0.6) for three measures (flexibility, integration, and recruitment) and four tasks (rest, Inscapes, movie, and flanker). For a given measure, global ICCs were highly similar across tasks (compare rows). For a given task, the locations of good ICCs were consistent across measures (compare columns). The peak ICC value was observed in the same location (γ=1.05, ω=2.05) in 7 out of the 12 two-dimensional γ-ω planes (highlighted by a black dashed line). The ICC score at this location was also good (>0.65) in the other 5 two-dimensional γ-ω planes. Thus, this parameter value pair was chosen as the optimal γ-ω values for our analyses. Note that the values in the parameter space where the number of communities was smaller than 2 or greater than 100 were set to zero in each plane. ICCs were evaluated with the maximal amount of data available (60 min).

**Figure 5.**
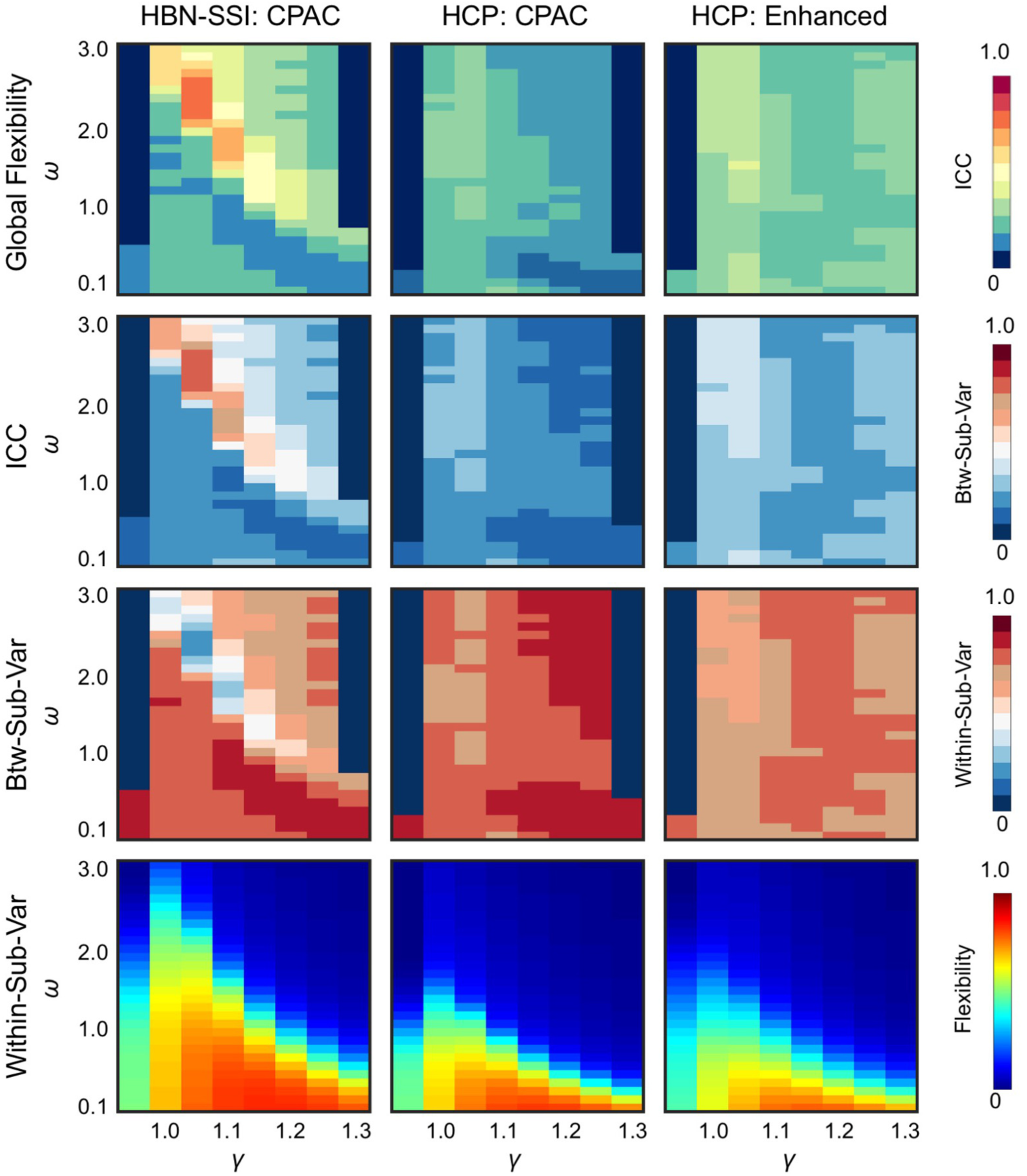
The γ−ω optimized for HBN-SSI cannot be generalized to HCP data. In HBN-SSI data, a range of parameters had good reliability (ICC≥0.6). However, in HCP data, we were unable to find a range of parameters with good ICCs regardless of preprocessing pipelines. As ICC was determined by both between-subject variance (Btw-Sub-Var) and within-subject variance (Within-Sub-Var), good ICCs in HBN-SSI overlapped with the portion of the landscape with high Btw-Sub-Var and low Within-Sub-Var. In HCP, the poor ICC was associated with low Btw-Sub-Var and high Within-Sub-Var. Furthermore, the two datasets also differed in global flexibility. The γ−ω optimized for HBN-SSI cannot be generalized to HCP data. In HBN-SSI data, a range of parameters had good reliability (ICC values; the values dropped more quickly when ω increased for the HCP data than for the HBN-SSI data. ICCs were estimated using 60 min of resting state data for test and retest. HBN-SSI: CPAC (HBN-SSI data preprocessed using CPAC pipeline) and HCP: CPAC (HCP data preprocessed using CPAC); HCP: Enhanced (HCP publically released extensively processed data).

### 3.2 Parameter optimization based on test-retest reliability

Because our goal is to optimize multilayer network-derived measures to study individual differences, we chose our parameters based on test-retest reliability scores. The parameter γ is the intra-layer coupling parameter, which defines how much weight we assign to the null network and controls the size of communities detected within a layer. The parameter ω is an inter-layer coupling parameter which defines the weight of the inter-slice edges that link a node to itself between two consecutive layers; it controls the number of communities formed across layers. We found that the selection of γ and ω had a large impact on the test-retest reliability of dynamic network measures for the HBN-SSI dataset (**Figure 4)**. Depending on parameter choice, test-retest reliability can range from poor to good. Overall, recruitment (mean ICC across the landscape: 0.54±0.11) is more reliable than integration (0.37±0.17), and integration is more reliable than flexibility (0.30±0.15). For each measure, the pattern of ICC values across the 2-dimensional parameter space is highly similar across tasks. For each task, the portions of the parameter space with good ICCs are consistent across measures. Thus, we were able to identify an optimal range of parameters generalizable across tasks and measures. For flexibility and integration, good ICCs (≥0.6) occur within a range of γ=[1.0-1.1] and ω=[1.7-3.0]. For recruitment, the range is broader: γ=[1.05-1.25] and ω=[1.2-3.0].

For the current analysis, we chose the parameters γ=1.05 and ω=2.05, which produce maximal ICC values in 7 of the 12 γ-ω planes and still produce relatively good ICC values (ICC>0.65) in the other 5 γ-ω planes. Turning to the parameter ω which affects coupling between layers, tuning it up to 2.05 yielded low estimates of flexibility. In a previous study, when the ω value was too high, flexibility values followed a heavy-tailed distribution with most values of flexibility equal to zero (i.e., close to a static network representation) (Telesford et al., 2016). In our investigation, the distribution of flexibility did not resemble this heavy-tailed distribution (**Figure S5A**), thus mitigating the potential concern that the parameter was tuned too high.

Because the ICC is determined by both within- and between-subject variability, good ICC could be caused by increased between-subject variability, decreased within-subject variability, or a combination of both. To understand the driver of this variation in test-retest reliability, we examined the landscape of dynamic measures, as well as the between- and within-subject variance of these dynamic measures. To make the variance values comparable, we normalized the between- and within-subject variance by the total variance. As expected, we found that the mean and variance of these dynamic measures also depended on the values chosen for γ and ω (**Figure S6**). The parameter values associated with good ICC overlapped with areas showing high between-subject variability and low within-subject variability, and largely overlapped with areas that had relatively low values of the dynamic measures (with a few exceptions for integration).

When the MPM was used, we found that reliability was poor for the previously recommended and commonly used values of γ=1 and ω=1. To better understand this poor reliability, we compared the recommended parameter choice with our reliability-optimized set. We found that although the spatial maps of flexibility were highly similar between two parameter choices (r=0.70), the magnitude of flexibility was much larger for [γ=1, ω=1] compared to [γ=1.05, ω=2.5]: 0.66±0.01 vs 0.16±0.01, respectively (**Figure S5A**). In the reviewed literature, when [γ=1, ω=1] was used, the range of flexibility is typically <0.25 (**Table 1**). This discrepancy is likely because previous studies used the MMM (Bassett et al. 2011, Bassett et al. 2013a, Bassett et al. 2013b, Bassett et al. 2015, Telesford et al. 2016, Finc et al. 2020). The poor ICC of [γ=1, ω=1] (mean: 0.19±0.21) relative to [γ=1.05, ω=2.5] (mean: 0.79±0.08) when the updated method was used was driven by the much lower between- and higher within-subject variance for [γ=1, ω=1] compared to [γ=1.05, ω=2.5] (except for the visual cortex).

To test whether functional parcellation and the resolution of a parcellation have an impact on dynamic reconfiguration, we repeated our analysis using Schaefer et al. (2018) 200 and 600 functional parcellations. We found that our results are relatively stable across different parcellations with the same number of ROIs, but parcellation resolution has an impact on parameter selection. Specifically, ICC values and dynamic network measures are similar between the Schaefer 200 and Craddock 200 atlases, and the optimized γ-ω are identical for the two atlases (γ=1.05 and ω=2.5) (**Figure S7**). However, when the number of ROIs increases from 200 to 600, values of dynamic reconfiguration measures increase while a higher γ value is required to achieve good ICC values (**Figure S8**). The optimized γ-ω for this higher resolution atlas is: γ=1.15 and ω=2.6. These results suggest that higher resolution atlases may augment values for node reconfiguration measures by increasing the number of communities. However, these measures of dynamic reconfiguration are less reliable as a higher inter-layer coupling parameter is needed to generate dynamic communities that span multiple layers. Moreover, higher values for the intra-layer coupling parameter are needed to reduce the number of communities to a level comparable to that of a low-resolution atlas.

To test whether the optimized parameters are generalizable, we applied the same multilayer analysis to HCP data and evaluated the test-retest reliability of flexibility. Compared to results for the HBN-SSI data, areas with relatively better reliability were located at values with lower γ and higher ω for the HCP data, although flexibility values were lower. Importantly, we were unable to identify any parameter value pairs with an ICC≥0.6 for the HCP data, and the overall reliability is poorer for the HCP data compared to HBN-SSI data (HBN-SSI mean: 0.30±0.15; HCP mean: 0.19±0.05) (**Figure 5**). This can be explained by lower between-subject variability and higher within-subject variability in the HCP data. To investigate the impact of preprocessing, we repeated our analysis on the HCP data using publicly released extensively preprocessed data. We found that both the flexibility values and reliability for the HCP data were similar between CPAC and HCP preprocessing, though the average reliability across the γ-ω landscape is higher for the HCP pipeline (CPAC mean: 0.19±0.05; HCP mean: 0.27±0.04). These results suggest that parameters optimized in one dataset (e.g., HBN-SSI, HCP) and/or preprocessing strategy (e.g., HCP pipeline employed ICA-FIX, while the C-PAC based pipeline used CompCor and GSR) may not be optimal for others (See **Figure 5** for visualization of γ-ω landscapes for HBN-SSI: CPAC, HCP: CPAC, HCP: Enhanced HCP).

Furthermore, we tested whether parameters optimized based on ICC are generalizable to parameters optimized for discriminability - a reliability index that is applicable to multivariate data (e.g., full brain or connectome). Similar to ICC, we found that discriminability was highest for recruitment, followed by integration, and lowest for flexibility (**Figure S9**). For recruitment, most discriminability values across the γ-ω plane were high (>0.9; Bridgford et al. 2019). For integration, high discriminability values were located in the portion of γ-ω with medium ICC. For flexibility, high discriminability values were located in the portion of γ-ω with small ICC. Our results illustrate that parameters optimized based on one criterion may not always be optimal for another criterion.

### 3.3 Data requirements for characterizing inter-individual differences in network dynamics

To establish the minimal data requirements for these types of analyses, we calculated the ICC for each measure and each task at six different scan durations: 10 min, 20 min, 30 min, 40 min, 50 min, and 60 min. Consistent with previous static analyses (Laumann et al. 2015, Xu et al. 2016, O’Connor et al. 2017), we found that the test-retest reliability of dynamic measures improves with increased scan duration, and that this pattern is consistent across tasks and across dynamic network measures (**Figure 6**). From 10 to 60 min, the largest improvement is from 10 to 20 min. After 40 min, most regions achieved good ICCs and improvements were less notable for longer scan durations. For regional and system-level variations in improvement of reliability as a function of scan duration, see **Figure S10**.

**Figure 6.**
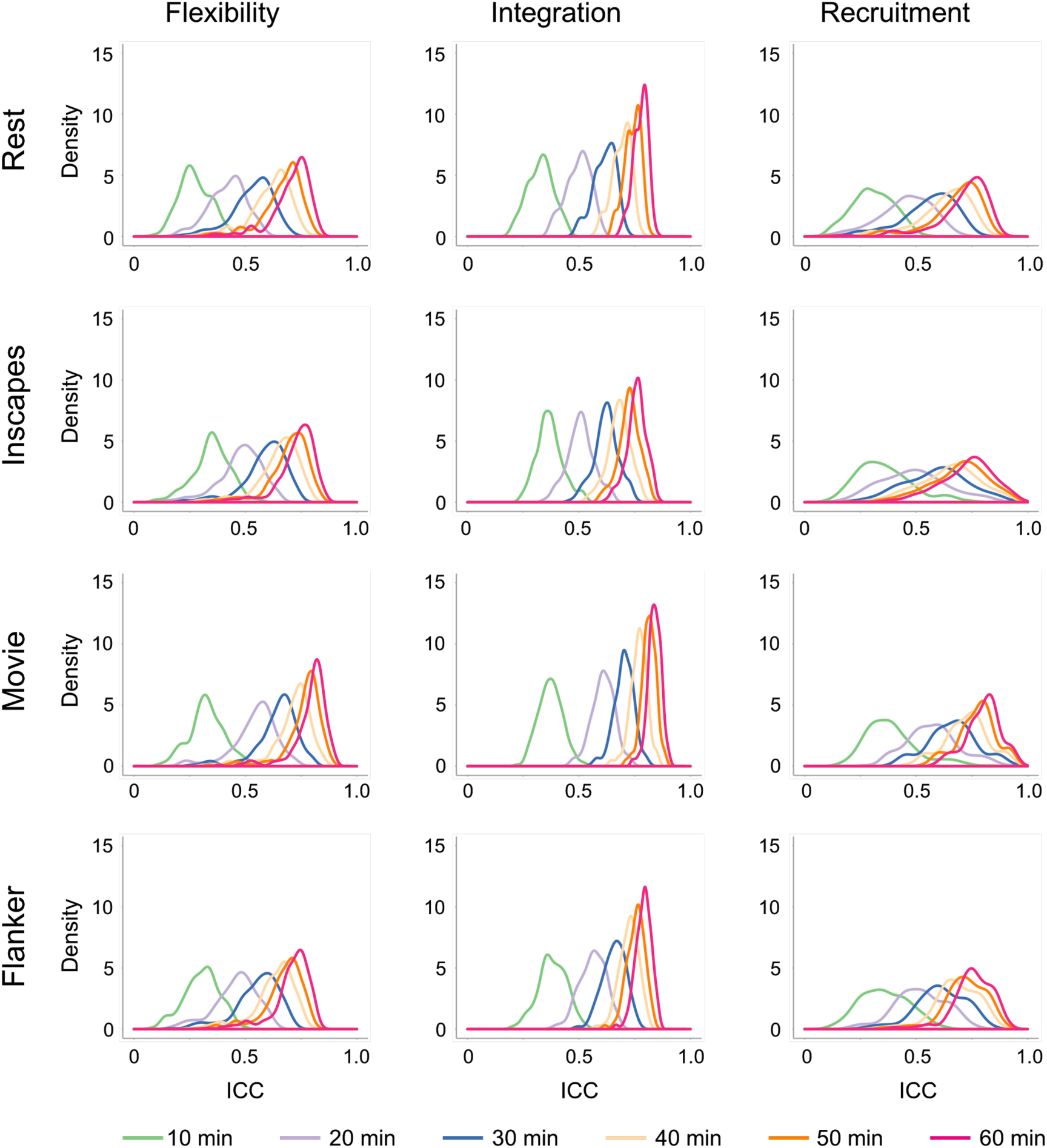
Test-retest reliability of dynamic network measures increases when the amount of data used for estimation increases. The density map of ICC values of 200 ROIs was plotted for three dynamic measures (flexibility, integration, and recruitment: columns) and four tasks (rest, Inscapes, movie, and flanker: rows) at six scan durations (10 min, 20 min, 30 min, 40 min, 50 min, and 60 min).

Regarding the question of how much data is needed for sufficient reliability, the answer depends on the criteria, the task, and the measure. Here, we define good test-retest reliability as over 50% of ROIs with ICC≥0.5 (Xu et al., 2016). For the movie condition, good test-retest reliability was achieved for all three measures at 20 min (81.5% of ROIs had an ICC≥0.5 on average across all three measures) (**Figure 7**). For the flanker condition, good reliability was achieved at 20 min for integration (83.0% of ROIs ICC≥0.5) and recruitment (57.0% of ROIs ICC≥0.5). For the rest and Inscapes conditions, good reliability was achieved at 20 min only for integration (52.0% and 55.5% of ROIs ICC≥0.5, respectively). With 30 min of data, all measures and all tasks had good test-retest reliability. Across scan duration and task condition, integration is more reliable than recruitment (Wilcoxon signed-rank test: p<0.001) and recruitment is more reliable than flexibility (p=0.02).

**Figure 7.**
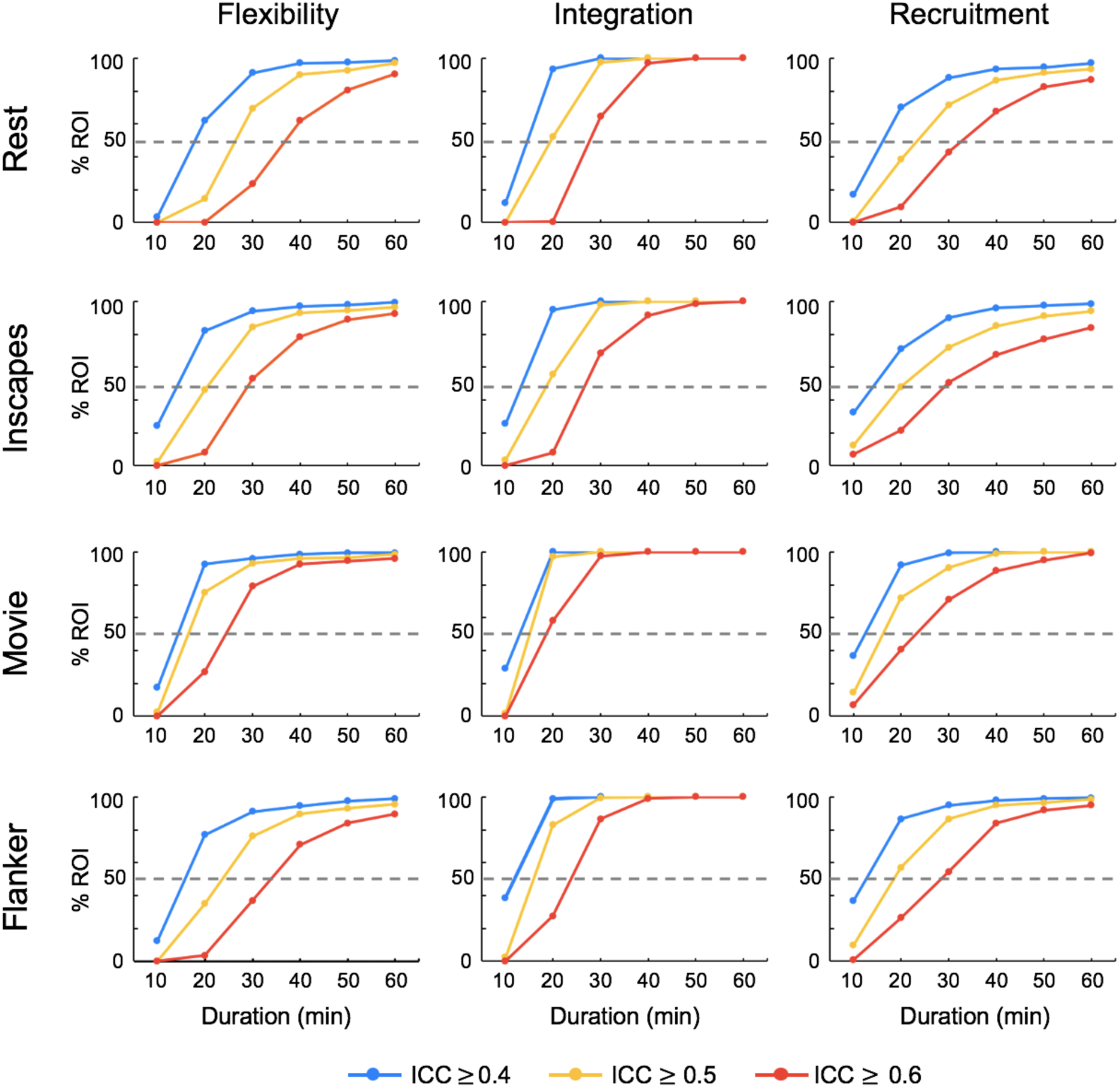
The minimal data requirements for sufficient reliability depending on the criteria, the measure, and the task. Percentage of ROIs with an ICC greater than 0.4 (blue line), 0.5 (orange line), and 0.6 (red line) were plotted for the three dynamic network measures (flexibility, integration, and recruitment: columns) and the four tasks (rest, Inscapes, movie, flanker: rows). The dashed grey line was drawn at 50%.

When data for one task is insufficient, a potential solution is to combine data across different tasks to increase scan duration, and thus improve reliability (O’Connor et al. 2017, Elliott et al. 2019a). To test whether this approach is relevant to the types of analyses performed here, we compared the ICCs obtained from 10 min of resting state data with the ICCs obtained from 10 min of Inscapes, movie, or flanker task condition, as well as those obtained from longer data created by adding either more resting state data or data from the other three tasks (**Figure 8)**. We found that at 10 min, ICCs are poor for all four tasks (over 75% of ROIs with ICC < 0.4), with resting state the poorest (96.5% of ROIs with ICC < 0.4). With increased scan duration, the reliability for the pure task condition increases, as shown in previous analyses, with the movie condition having the most ROIs showing good ICC at each scan duration. When comparing pure resting state data with mixed data of the same duration, generally the pure data had a greater number of ROIs with fair or good to excellent ICC compared to the mixed data. When comparing 10 min of resting state data with longer mixed data, we found that combining data from different tasks improved reliability. The degree of improvement depended more on how much data was combined, and less on what task conditions were combined.

**Figure 8.**
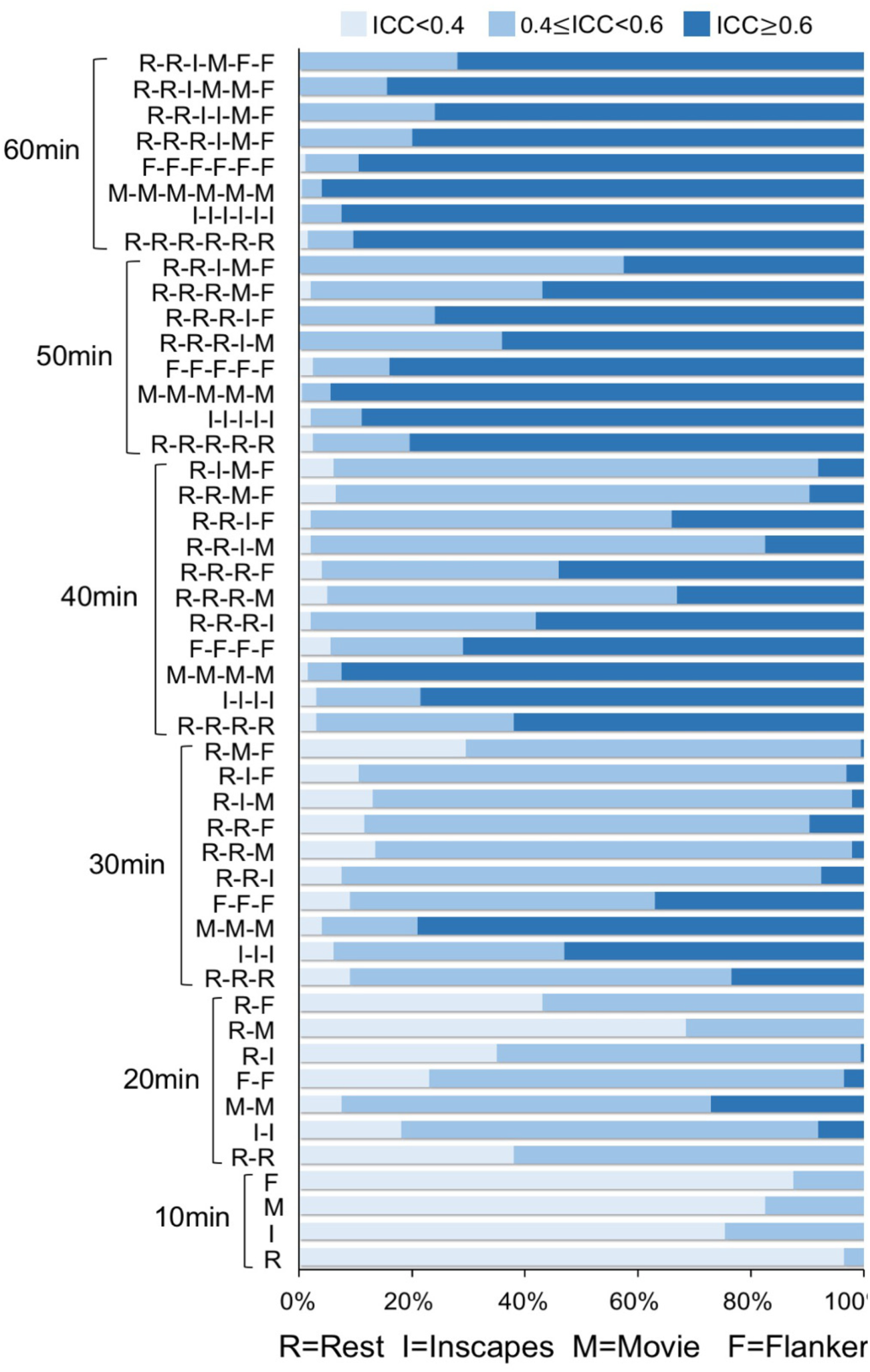
Combining data from different tasks improved reliability. Percent of ROIs showing poor (light blue: ICC<0.4), fair (medium blue: 0.4≤ICC<0.6), or good (dark blue: ICC≥0.6) reliability were plotted for six durations: 10min, 20 min, 30 min, 40 min, 50 min, and 60 min. For each duration, the data can be a single condition or a combination of the four conditions: rest (R), Inscapes (I), movie (M), and flanker (F). Each letter (the abbreviation of each condition) represents 10 min of data.

### 3.4 Task modulation on test-retest reliability of network dynamics: hierarchical linear mixed model

To separate variation among scan conditions from variations between sessions, we assessed between-condition reliability and between-session reliability simultaneously in a hierarchical linear mixed model. With the optimized γ-ω and the maximal amount of data available (60 min), we found that both between-session (two sessions, 60 min/session) and between-condition (four conditions) reliability were excellent (between-session median ± interquartile range: flexibility, 0.76±0.05; integration, 0.80±0.02; recruitment, 0.77±0.08; between-condition: flexibility, 0.74±0.10; integration, 0.76±0.07; recruitment, 0.77±0.16) (**Figure 9**). Consistent with previous work (O’Connor et al., 2017), we found that between-condition reliability of the visual and somatomotor network tended to be the poorest for recruitment which quantifies within-network functional interactions. Because different task states vary systematically in the richness of visual stimuli (movie>Inscapes>flanker>rest) and motor demands (flanker>the other three conditions), it is reasonable that these primary networks re-configure themselves according to unique task demands.

**Figure 9.**
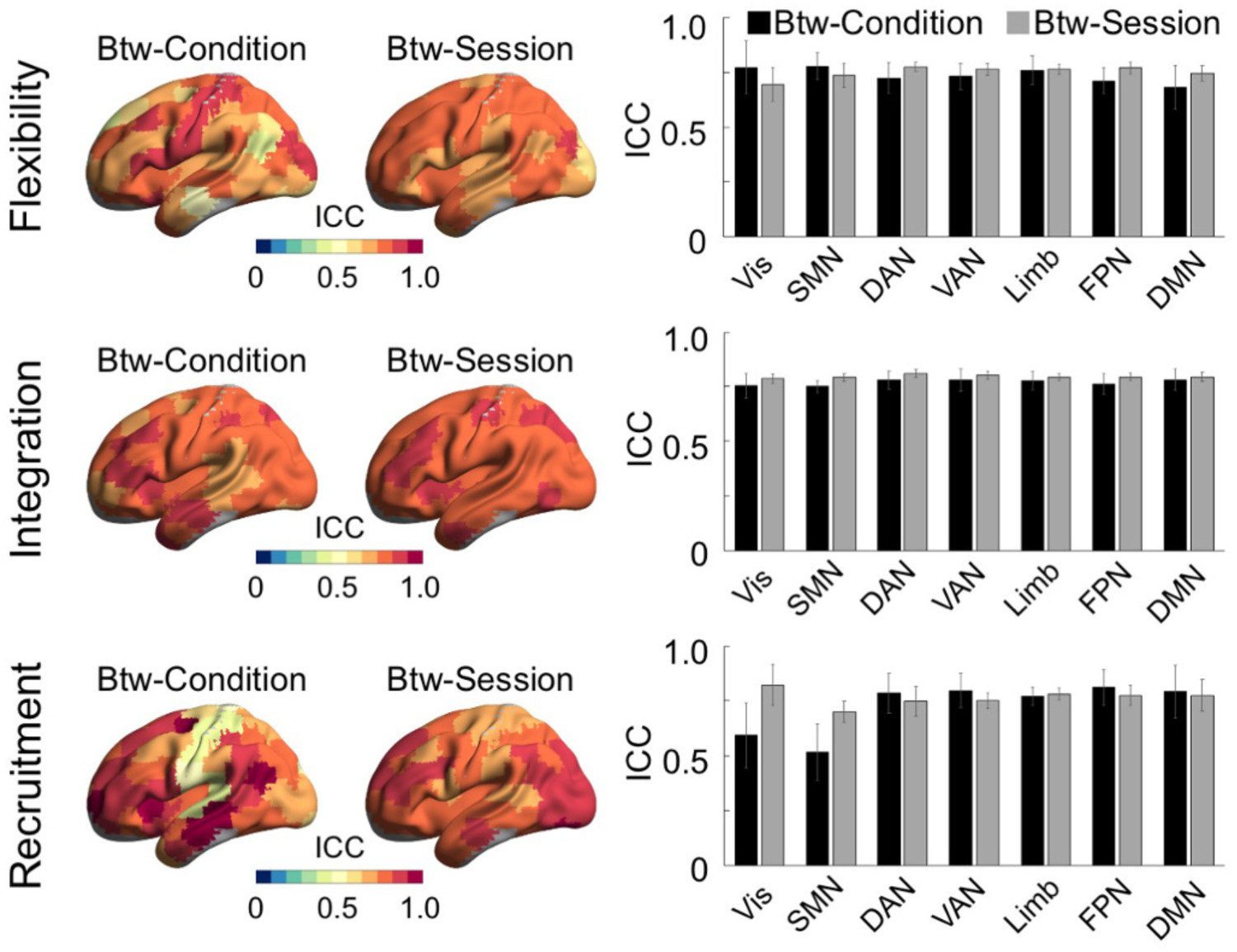
Both between-session and between-condition reliability evaluated in a hierarchical linear mixed model were good to excellent for 60 min of data. The between-condition (btw-condition: reliability between rest, Inscapes, movie, and flanker) and between-session (btw-session: reliability between test and retest) ICCs were plotted on the surface map using BrainNet Viewer (Xia et al. 2013), as well as summarized per the seven networks defined by Yeo et al. (2011) in bar plots. Rows: flexibility, integration, recruitment. Vis: visual network; SMN: somatomotor network; DAN: dorsal attention network; VAN: ventral attention network; Limb: limbic network; FPN: frontoparietal network: DMN: default mode network. The same network abbreviations were used for subsequent figures.

### 3.5 Task modulation on test-retest reliability of network dynamics: linear mixed model

Following the high-level model, we investigated test-retest reliability for each task separately using simple linear mixed models. We found that all four tasks have good to excellent test-retest reliability for all three measures (**Figure 10**). Median ± interquartile range of ICC for rest, Inscapes, movie, and flanker were: flexibility (0.73±0.09, 0.75±0.09, 0.81±0.07, 0.73±0.08), integration (0.78±0.05, 0.76±0.05, 0.84±0.04, 0.79±0.05), and recruitment (0.74±0.13, 0.74±0.16, 0.81±0.09, 0.76±0.11). When reliability was directly compared between tasks, there was a significant main effect of task for all three measures (Friedman test: p<0.001). Using *post hoc* testing, we found that the movie condition displayed significantly better test-retest reliability in all dynamic network measures than the other three conditions (Wilcoxon signed-rank test: all p-values<0.001, below Bonferroni correction for 18 tests: 3 measures×6 possible pairing).

**Figure 10.**
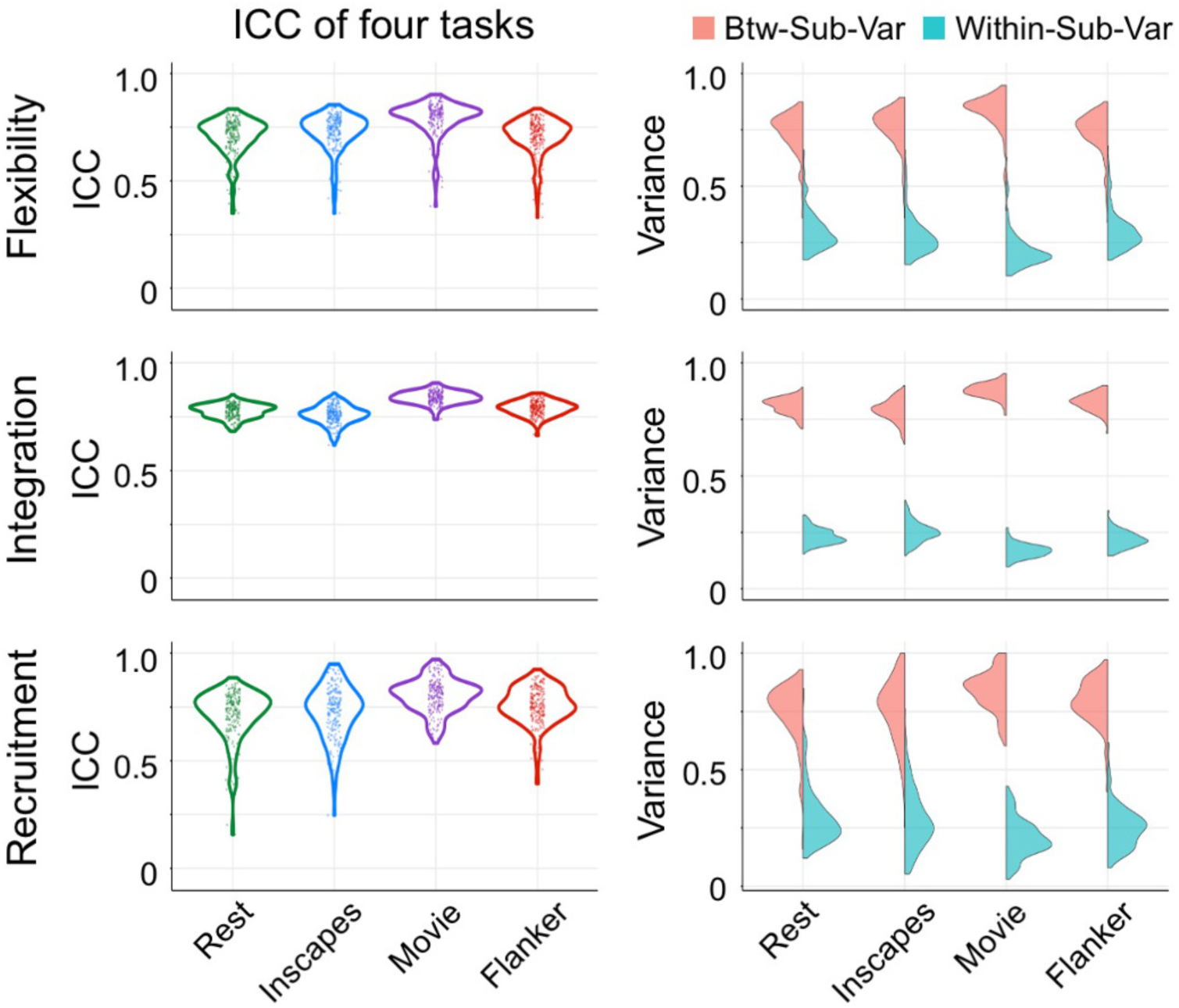
The movie condition was most reliable. Distribution of ICCs of 200 ROIs were plotted for four conditions (rest, Inscapes, movie, and flanker) in the left column. Density of between-subject variance (Btw-Sub-Var: salmon) and within-subject variance (Within-Sub-Var: light sea green) were plotted for each condition in the right column. Rows: flexibility,

For the comparison of the remaining conditions, the results were measure dependent. For flexibility, test-retest reliability in the Inscapes condition was significantly higher than in the flanker condition (p<0.001, corrected), and the other comparisons were not significant; for integration, reliability differed significantly (flanker>rest>Inscapes, p<0.001, corrected); for recruitment, reliability in the flanker condition was also significantly higher than in the rest and Inscapes conditions (p<0.001, corrected). Generally, the reliability of these dynamic measures did not simply increase as a function of task engagement. Higher ICC scores were typically associated with relatively higher between-subject variance and lower within-subject variance (**Figure 10**).

After considering overall reliability (median ICC), we next visualized regional and network differences in reliability between tasks. Consistent with overall results, we found that the movie condition exhibited higher reliability than the other three conditions in most brain regions and networks (**Figure 11**). The other three conditions are similar to each other with a few exceptions: flexibility of the somatomotor, visual, and default mode networks, and recruitment of the visual and somatomotor networks. The observation that task effects were most robust within the primary cortices is consistent with the hierarchical linear mixed model and with previous work (O’Connor et al., 2017). Furthermore, we found that the spatial topographies of ICC differ among dynamic measures. This is expected because different measures capture different features of the dynamic community structure. Overall, the spatial variation of ICC was small for integration at 60 min (i.e., all regions showed good to excellent reliability) across conditions. The spatial topography for flexibility and recruitment seems more complex. For example, the visual cortex was most reliable for recruitment during movie conditions but least reliable for flexibility. These results may suggest that the likelihood that a visual region will be assigned to the same community as other visual network regions (quantified by recruitment) is stable during the movie watching condition when visual stimuli were continuously presented. However, the number of times that a visual region switches its community membership (captured by flexibility) is not as reliable.

**Figure 11.**
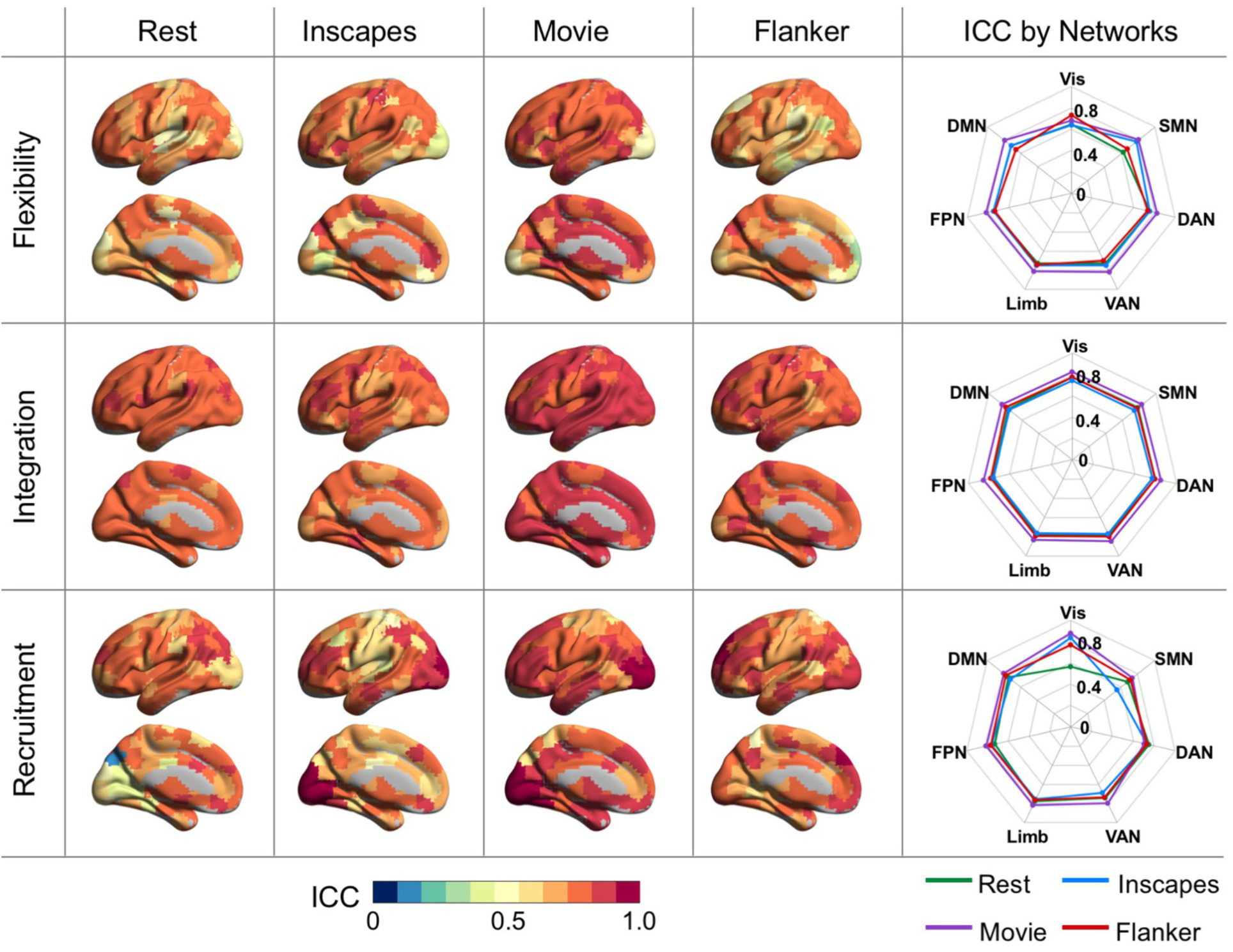
The impact of condition on the test-retest reliability of dynamic network measures. Spatial maps of ICCs for rest, Inscapes, movie, and flanker condition are shown on the brain surface for flexibility, integration, and recruitment. ICCs of 200 ROIs were averaged based on Yeo et al. (2011)’s seven networks for each of the four conditions and shown in the radar chart: Rest (green line), Inscapes (light blue line), Movie (purple line), and Flanker (red line).

### 3.6 Addressing concerns regarding head motion

Head motion remains a major concern for dynamic functional connectivity estimation (Yang et al. 2014, Bassett et al. 2018, Satterthwaite et al. 2019). In the present work, we only included participants with minimal head motion. During preprocessing, we regressed out 24 motion-related parameters (Friston et al. 1996); as well as controlled motion with more generalized approaches such as global signal regression at the individual level (Yan et al. 2013, Yang et al. 2014, Lydon-Staley et al. 2019a). To provide further insights into this concern, we examined the correlation with head motion, which we quantified as median framewise displacement (Jenkinson et al. 2002), and the global mean of each dynamic measure; we did not observe any significant correlations between these variables. Furthermore, we re-estimated test-retest reliability for flexibility on the movie condition using the optimized parameter while including median framewise displacement as a covariate at the group level in the linear mixed model. We found similarly good reliability with and without head motion included in the model (ICC=0.67 and 0.74, respectively), suggesting that the impact of head motion on test-retest reliability was small.

## 4 Discussion

Optimizing the reliability of dynamic network methods is key to accurately characterizing trait-like individual differences in brain function. The present work examined the impact of the modularity maximization algorithm, network parameter selection, scan duration, and task condition on the test-retest reliability of dynamic network measures obtained using multilayer network models. We found that each of these factors impacted reliability to differing degrees. As suggested by prior work, optimal parameter selection was found to be an important determinant of reliability; interestingly, our findings revealed a more complex story than previously appreciated, as reliability across the multivariate parameter space was found to depend on an update in the multilayer community detection algorithm. Consistent with findings from the static functional connectivity literature, scan duration was found to be a much stronger determinant of reliability than scan condition. As discussed in greater detail in the following sections, our findings suggest that optimization of multilayer network models per dataset is essential rather than selecting a single set of parameters or methods previously used in the literature.

Multilayer network models of neural systems offer the potential to illuminate time-varying aspects of brain function that could not otherwise be revealed from traditional static network approaches. The aim of the present work is to quantify and optimize test-retest reliability for multilayer network models, establish minimum data requirements for obtaining reliable dynamic measures, and identify which task condition(s) can provide a reliable context in which to investigate time-invariant network dynamics. Although we only evaluated the reliability of the GenLouvain algorithm, which is one of the most popular algorithms, the framework presented can be applied to other software and multilayer network modularity-maximizing algorithms. With increased interest in linking the time-resolved reconfiguration of functional brain networks to normal cognition and disorders, it is important for the field to establish standards to guide the application of dynamic connectivity approaches. The present work systematically evaluated the impact of parameter selection, scan duration, and task condition on the test-retest reliability of dynamic network measures, which addresses an important gap in the literature. Although the present work focused on fMRI data, our analytical methods and results are broadly applicable, as multilayer network modeling has been widely applied to other neuroimaging modalities (e.g., structural MRI, EEG, MEG) and non-neuroimaging data, such as social, economic, gene, and protein networks (Boccaletti et al. 2014).

### 4.1 A cautionary note on the selection of GenLouvain algorithms

When using the generalized Louvain algorithm for modularity maximization, a critical step is to merge nodes into communities. The MMM is the default method used in the implementation of this algorithm, which merges nodes that produce the highest increase in modularity. In 2016, a newer approach was introduced, the MPM, which selects merges based on the probability distribution of modularity increases. A previous study of financial data reported that when the default method was used, two computational issues arise in the multilayer setting: an under-emphasis of persistence and an abrupt drop in the number of intra-layer merges in certain portions of the parameter space - both of which can lead to an abrupt change in the quantitative measure derived (Bazzi et al. 2016). Here, when the default method was used, we observed an abrupt drop-off in the optimization landscape in brain imaging data as well as in synthesized data (**Figure 3**).

The abrupt discontinuity in the γ-ω landscape is not a coding error. Instead, it is a reflection of how the edges are defined (e.g., functional neuroimaging datasets generally use normalized correlation/coherence values between regions/voxels), the relationship between the multilayer inherited parameters, and the strengths of edges. The abrupt change occurs when the inter-layer coupling parameter (ω) is greater than the average intra-layer edge strength. As the optimization of the modularity function has proven to be non-deterministic polynomial-time-hard (Newman 2006), the default deterministic algorithm suffers from local minimum issues and results in fewer merges - especially when ω values are high. In contrast, the probabilistic approach, with a mechanism analogous to a simulated annealing algorithm, is better suited to approximate the global optimum of the quality function (Bazzi et al., 2016). Using the probabilistic approach leads to more variability across multiple simulations for a given dataset, thus mitigating the abrupt change seen in the optimization landscape.

The fact that abrupt discontinuities were observed consistently regardless of data type raises concerns regarding the accuracy of dynamic measures derived using the default option and with parameters selected above the point of apparent discontinuity in the 2-dimensional parameter space. Based on these results, as well as our demonstration that the Modularity Probability Method has higher test-retest reliability and better validity compared to the Maximum Modularity Method, we strongly recommend that investigators use the updated method for multilayer network analysis, especially when applied to ordinal or temporal networks. Caution should be taken when attempting to judge findings obtained with the MPM based on older findings obtained with the MMM.

### 4.2 Parameter optimization for multilayer network analyses

To detect community structure, we employed the most commonly used algorithm to maximize the multilayer modularity quality function (Mucha et al. 2010). Communities that are detected using this algorithm are highly dependent on free parameters (i.e., γ and ω), thus we aimed to explore the space defined by these parameters and identify optimal parameter selection ranges in terms of test-retest reliability. As one parameter may affect the other parameter’s optimal setting, it can prove useful to optimize γ-ω jointly. Although several heuristics exist for choosing the “best” value of γ and ω (Bassett et al. 2013a, Chai et al. 2016, Weir et al. 2017), optimizing the ICC has not previously been proposed, possibly because it requires the acquisition of a retest dataset. Our results suggest that a systematic evaluation of the parameters in terms of reliability has marked utility, as parameter choices directly impact reliability.

In the 2-dimensional parameter space of the γ-ω plane, we were able to find a range of parameters that produced dynamic network measures of community structure in multilayer networks with good reliability. For flexibility and integration, betterer reliability was achieved with higher ω (i.e., when there is a strong temporal coupling) and lower γ (i.e., when there are fewer communities). For recruitment, good reliability was achieved with high ω and a wide range of γ from low to high. Stronger temporal coupling in a multilayer network is typically associated with lower temporal variability in network partitions over time. The good test-retest reliability obtained at high ω and low γ, for flexibility and integration, may suggest that the temporal variability reserved after tuning up ω is composed of more between-subject variability than within-subject variability when the number of communities is small. The relative insensitivity of recruitment to the number of communities may be explained by our choice of predefined systems in which ROIs tend to be grouped together over time. These results suggest that ICC-guided parameter selection can potentially maximize between-subject variability and minimize within-subject variability. This practice is consistent with the recent call for including assessment and optimization for reliability as a common practice in neuroimaging, as it helps to improve statistical power and decrease the amount of data required per subject (Zuo et al. 2019).

A critical cautionary note for the identification of an optimal parameter set comes from the dependence of reliability across the 2-dimensional parameter space on the specific modularity maximization algorithm. A parameter choice of [γ=1, ω=1] was recommended in the literature based on modularity and partition similarity, as well as the differences between measures estimated on a real network compared to an appropriate multilayer network null model (Bassett et al., 2013a). Following these initial publications (Bassett et al., 2011, Bassett et al., 2013a), most studies have used [γ=1, ω=1] as their parameter choices (see **Table 1**) and tested the robustness of this parameter selection with small variations. Given this parameter choice falls in the dropoff area when the default MMM was used (**Figure 3 and Figure S1)** and it also falls in the poor test-retest reliability area when the updated MPM was used (**Figure S5**), the parameter choice of [γ=1, ω=1] needs to be reconsidered.

### 4.3 Generalizability of the optimized parameters to HCP data

To determine whether the parameters optimized for one dataset can be generalized to a different dataset, we compared the reliability landscapes from the HBN-SSI to those obtained with the HCP Test-Retest dataset. Recognizing the differences in preprocessing, we compared two datasets with identical preprocessing using CPAC. To investigate the impact of preprocessing, we also repeated the HCP analysis using the publicly released preprocessed data using the Enhanced HCP pipeline. We found that the reliability between the HCP Test (60 min) and Retest (60 min) data was much lower than that observed in the HBN-SSI dataset across the 2-dimensional γ-ω parameter space and this pattern was consistent regardless of preprocessing pipelines. One potential explanation for this difference between two datasets is that HCP data were acquired using faster sampling than the HBN-SSI data (TR: 0.72 s vs. 1.45 s). While static studies have indicated that increasing temporal resolution can either improve (Birn et al. 2013, Liao et al. 2013, Zuo et al. 2013) or have no impact (Horien et al. 2018) on reliability, the opposite was observed for dynamic analyses (Choe et al. 2017). In addition to TR, these two datasets were collected in scanners with different magnet strength (1.5 T vs 3 T) and used different multiband factors (3 vs 8 for HBN-SSI and HCP, respectively). Furthermore, although the subjects in the HCP and HBN-SSI were similar in terms of participant age and sex (HBN-SSI: 29.8±5.3 years old, 50% males; HCP: 30.32±3.28 years old, 36% males), intervals between acquistion of test and retest datasets differed. For the HBN-SSI, the sessions were acquired within two months, while the HCP has a large variation in test-retest interval (range from 52 to 326 days: 133.36±58.27 days). These factors may impact the reliability of network flexibility assessment.

Importantly, our results raise significant concerns about the potential dependencies of ‘optimal parameters’ for multilayer network analysis on the datasets employed, beyond the specific properties identified in our work (i.e., amount of data per subject, or per condition). Demonstrating that test-retest reliability can differ substantially between datasets is a significant concern, as it suggests that parameters optimized in one dataset may not be optimal for others. Compounding the challenge at hand, few data collection efforts include test-retest samples, and few contain the amounts of data per subject that our analyses suggest may be needed to achieve sufficient reliability. These varying factors raise concerns about the appropriateness of applying this approach to datasets that do not have a retest sample. It is also important for future studies to assess the generalizability of parameter optimization to datasets harmonized for key aspects of undesirable non-biological sources of variation, such as scanner manufacturer, acquisition protocol, and preprocessing steps. If such datasets are not available, applying statistical harmonization techniques, such as ComBat (i.e., combining batches) (Johnson et al. 2007, Fortin et al. 2018, Yu et al. 2018), may decrease unwanted site effects and optimize multilayer network analysis.

### 4.4 Minimal data requirements for obtaining reliable dynamic estimates

Many factors impact the test-retest reliability of functional connectivity-based measures, among which scan duration is one of the most important (Zuo and Xing 2014, Zuo et al. 2019). Establishing minimal data requirements to obtain reliable estimates is an active research area for static connectivity analysis (Van Dijk et al. 2010, Anderson et al. 2011, Birn et al. 2013, Liao et al. 2013, Zuo et al. 2013, Laumann et al. 2015, Xu et al. 2016, Noble et al. 2017, Tomasi et al. 2017). However, to date, few efforts have been made to determine the scan duration needed to obtain reliable estimates of dynamic network measures. Here, we found that the test-retest reliability of dynamic network measures was poor for 10 min of data; it improved greatly when data increased to 20 min for movie fMRI and to 30 min for the other scan conditions. While increased scan duration has consistently been shown to improve reliability, studies vary in conclusions about the necessary data required to obtain reliable estimates. Studies have suggested that 5-10 min of data are sufficient to achieve respectable test-retest reliability (Van Dijk et al. 2010, Liao et al. 2013, Zuo et al. 2013, Tomasi et al. 2017); importantly, these studies have either focused on the default and frontoparietal networks, which have better reliabilities than other functional networks, or used more complex derived measures than simple edgewise complexity. More recent work has convergently reported a substantial improvement in reliability to a level more useful for characterizing trait-like individual differences when data are increased from 5-10 min to 20-30 min (Laumann et al. 2015, Xu et al. 2016, Noble et al. 2017, O’Connor et al. 2017, Elliott et al. 2019a). Our results are consistent with these static functional connectivity studies.

As temporal dynamic analyses are susceptible to spurious variations (Hutchison et al. 2013, Leonardi and Van De Ville 2015, Lehmann et al. 2017), one would assume more data are required to obtain reliable measures for dynamic analyses compared to static analyses. Instead, our data recommendations for estimating flexibility, recruitment, and integration from multilayer community detection analyses to examine trait-like individual differences are comparable to those for static functional connectivity analysis (20∼30 min). This result may reflect our having optimized the analyses for test-retest reliability. As previous multilayer network-based studies vary widely in scan duration (ranging from 5 min to 3.45 hours: **Table 1**), it is crucial to establish minimal data requirements for the study of trait-like individual differences.

### 4.5 Improvement of test-retest reliability by combining different conditions

It may not be practical to collect 20 to 30 min of data for a single condition, which motivates the question of whether different conditions can be combined to increase scan duration and improve test-retest reliability. Our hierarchical linear mixed model revealed good between-condition reliability, as well as good between-session reliability. These results are consistent with previous static connectivity analysis using the HBN-SSI dataset which demonstrated good between-condition reliability (O’Connor et al., 2017). Our findings are also consistent with previous work showing that task and resting-state data share a large proportion of variance (Cole et al. 2014, Geerligs et al. 2015) and that inter-task variance is much smaller relative to inter-subject variance in functional connectivity (Finn et al. 2015, Gratton et al. 2018). Recent work leveraging shared features across resting-state and task fMRI using a method called ‘general functional connectivity’ has demonstrated that intrinsic connectivity estimated based on a combination of task and resting-state data offers better test-retest reliability than that estimated from the same amount of resting state data alone (Elliott et al., 2019a). Here, we also found that when scan duration was increased from 10 to 20, 30, or 40 min by combining task and resting-state data, the reliability of flexibility was greatly improved to a degree comparable to that estimated from 20, 30, or 40 min of resting data alone. Extending our understanding beyond prior studies of static connectivity, our results suggest dynamic network reconfiguration is similar across conditions when scan parameters and duration are optimized, thus supporting the feasibility of combining data from different tasks and conditions to improve reliability.

### 4.6 Movie fMRI identified as the most reliable condition

Another factor that impacts test-retest reliability of brain imaging-based measures is experimental paradigm due to the condition-dependent nature of brain activities (Zuo, Xu, & Milham, 2019). Multilayer networks have been used to assess network reconfiguration during resting state (Mattar et al. 2015, Betzel et al. 2017, Wei et al. 2017, He et al. 2018, Khambhati et al. 2018, Pedersen et al. 2018, Zheng et al. 2018, Al-Sharoa et al. 2019, Feng et al. 2019, He et al. 2019, Li et al. 2019, Shao et al. 2019, Tian et al. 2019, Lydon-Staley et al. 2019a, Lydon-Staley et al. 2019b), as well as during controlled cognitive tasks (Bassett et al. 2011, Bassett et al. 2015, Braun et al. 2015, Chai et al. 2016, Telesford et al. 2016, Schlesinger et al. 2017a, Schlesinger et al. 2017b, Gerraty et al. 2018, Cooper et al. 2019). The present work extended previous studies by including naturalistic viewing paradigms. Naturalistic paradigms offer increased ecological validity and allow researchers to study highly interactive dynamic cognitive processes (Bottenhorn et al. 2019) and probe complex multimodal integration (Sonkusare et al. 2019). Thus, studies characterizing network dynamics and establishing test-retest reliability of these paradigms together have the potential to enhance our understanding of cognition as it occurs more naturally. A recent meta-analysis revealed that naturalistic paradigms recruit a common set of networks that allow separate processing of different streams of information as well as integration of relevant information to enable flexible cognitive and complex behavior (Bottenhorn et al., 2019).

Compared to a passive resting state and an active flanker condition, we found the movie condition had the best test-retest reliability. These results are consistent with previous static network studies which suggested better test-retest reliability for movie conditions when compared to resting state (Wang et al. 2017). Naturalistic viewing was shown to have enhanced ability to identify brain-behavioral correlations compared to conventional tasks (Cantlon and Li 2013, Vanderwal et al. 2019) and was less impacted by head motion (Vanderwal et al. 2015), especially for pediatric samples. Some have suggested that the better reliability may be explained by the enhanced ability of movie watching to detect inter-individual differences in functional connectivity that are unique at the individual level compared to resting state (Vanderwal et al. 2017); alternatively, findings might be related to the increased level of engagement for movies, which is known from the task fMRI to help stabilize connectivity patterns over time (Elton and Gao 2015). Regardless of explanation, the present results support the utility of naturalistic paradigms for investigating network dynamics in developmental and clinical applications.

### 4.7 Limitations and future work

To estimate functional connectivity, we used wavelet coherence based on its predominance across similar studies in the literature (see **Table 1**), as well as due to its advantages in terms of denoising, robustness to outliers, and appropriateness for fMRI time series (Zhang et al. 2016). While wavelet coherence offers several advantages, it is a frequency-specific measure and does not utilize phase information (Percival and Walden 2000). As such, wavelet coherence is not useful when the phase of the signal is critical. Ongoing work is examining the reliability of other connectivity estimation methods, such as the Pearson’s correlation coefficient (Bassett et al. 2011, Mattar et al. 2015, Chai et al. 2016, Pedersen et al. 2018), which is informed by both phase and frequency information and which can be computed more swiftly. Future work should investigate how edge density and threshold as well as edge weight sign (i.e., inclusion/exclusion of negative correlations) might impact the reliability of the dynamic network measures studied here.

We focused our analyses on low frequency fluctuations (0.01-0.1Hz). The poorer reliability of the flanker condition compared to the movie condition could reflect the fact that we ignored the high frequency signals in the flanker task. To evaluate this possibility, we assessed flanker data reliability at a higher frequency range: 0.1-0.3 Hz. This range was selected to avoid the noisy upper bound (with TR=1.45 s, the highest frequency we can examine is 0.34 Hz). We found that the reliability of dynamic measures obtained in the low frequency signals of the flanker task was much higher than in the higher frequency signals (**Figure S11**). This suggests that the low frequency signals carry more non-random between-subject variation for this task, and that the relatively poor reliability of the flanker condition compared to the movie condition cannot be explained by frequency alone. Alternatively, the poorer reliability of the flanker condition could be ascribed to its having been designed to minimize between-subject variance to “isolate” a single cognitive process (Elliott et al. 2019b).

We determined the size of the parameter space by considering the number of communities (≥2 and ≤100), and we estimated the ICC at each point in the 2-dimensional γ-ω parameter space at a relatively coarse scale (γ: 0.9-1.3 with increments of 0.05; ω: 0.1-3.0 with increments of 0.1). We note that this resolution is comparable to most previous work (Bassett et al. 2011, Bassett et al. 2013b, Braun et al. 2015, Braun et al. 2016, Chai et al. 2016, He et al. 2018). Recent extensions of the multilayer network approach to dynamic community detection have demonstrated that sweeping across a range of intra-coupling parameters can offer insights into the multi-scale hierarchical organization of the brain (Ashourvan et al. 2019). Moreover, such studies have demonstrated that inter- and intra-subject variability in modular structure are scale specific (Betzel et al. 2019). Thus, sampling community structure from more points in the γ, ω parameter space may provide a better characterization of the brain’s dynamic network reconfiguration.

Indeed, some algorithms have been developed recently which allow a more refined and efficient search for parameters, for example, the Convex Hull of Admissible Modularity Partitions (CHAMP) (Weir et al. 2017). Unlike the traditional way of selecting parameters in which the optimal partitions obtained at each (γ, ω) were treated independently, CHAMP uses the union of all computed partitions to identify the convex hull of a set of linear subspaces. It can greatly reduce the number of partitions that can be considered for future analyses by eliminating all partitions that were suboptimal across a given range of parameter space. Although the CHAMP software package is currently in its early versions (https://github.com/wweir827/CHAMP), future work implementing these methodological updates can potentially facilitate the parameter optimization process and map the ICC landscape in greater detail.

We found that our parameter selection was stable across functional parcellations with the same resolution. However, it was sensitive to the resolution of a parcellation (i.e., number of ROIs of a parcellation). Recent work further demonstrated that functional parcel definitions change with task (Salehi et al. 2020a) and individualized functional networks reconfigure with cognitive state (Salehi et al. 2020b). Thus, another limitation of the present work is that we used fixed nodes and did not consider flexible functional nodes. It is important for future work to evaluate the test-retest reliability of multilayer network measures computed using flexible functional nodes and taking into consideration the resolution of a parcellation. As the network measures we computed are summary measures of dynamic reconfiguration, another limitation is that they did not have the temporal resolution to relate to changing conditions in the movie or flanker task.

A further limitation is that we optimized parameters based on the global mean of dynamic network measures computed across the whole brain. It is possible that each network may have different optimal parameters and the parameters optimized at the global level may not be optimal at the network level. It is important for future work to test this possibility and extend the current maximization framework further. Additionally, we fixed the values of γ and ω to be uniform across all layers as done in prior work (**Table 1**). Another extension of the present investigation is to devise heuristics for determining the values of these parameters in a layer-specific way, allowing for finer control over the features of detected communities.

Optimization of multilayer network measures for reliability has the potential to enhance our ability to use these measures and study trait-like brain-behavior relationships more efficiently(Choe et al. 2017, Zuo et al. 2019). Establishing good reliability is a key component of reproducible research (Nichols et al. 2017, Poldrack et al. 2017). However, good test-retest reliability does not necessarily correspond to high sensitivity to detect brain-behavior relationships (Noble et al., 2017). Thus, it is important for future work to investigate the functional relevance of reliability-optimized dynamic network measures, as well as to consider optimizing the multilayer modularity framework based on other factors, such as predictive accuracy (Dadi et al. 2019). Prior work suggests that pipelines optimized on discriminability can better detect brain-phenotypic associations (Bridgford et al. 2019). Other prior work suggests that pipelines optimized on predictive accuracy give the best prediction for diverse targets (including neurodegenerative diseases, neuropsychiatric diseases, drug impact, and psychological traits) across multiple datasets (Dadi et al., 2019). Thus, adding these new dimensions as optimization targets may enhance the ability of multilayer network measures to become fundamental tools to delineate meaningful brain-behavior relationships. This approach may be particularly useful for examining developmental questions. Multi-layer network analyses have been applied to reveal developmental patterns in brain function (Betzel et al. 2015, Schlesinger et al. 2017b). Changes in brain connectivity dynamics have also been reported in the context of other dynamic connectivity methods from childhood to adulthood (Faghiri et al. 2018, Vohryzek et al. 2019), during adolescence (Medaglia et al. 2018), and across the lifespan (Yan et al. 2017).

## 5. Conclusions

The application of dynamic (i.e., time-varying) graph measures to fMRI data is a rapidly growing area and there is a clear need in the network neuroscience field for reliable measures that can be used to find trait-like individual differences in cognition and disorders. Our results provide evidence that dynamic measures from the well-known multilayer community detection technique (multilayer modularity maximization) can be reliable when the updated multilayer community detection method is used, the parameters are optimized for reliability, and scan duration is sufficient. However, we do not assert that our results are directly applicable to any other dataset. Instead, we highlight concerns about generalizability arising from our difficulties finding a robust optimization that would generalize across the datasets tested. Our results caution the field that continued optimization of multilayer network models is needed before any single set of parameters or methods can be accepted as standard practice. Future work is needed to continue optimizing this framework by evaluating the impact of imaging parameters (e.g., sampling rate), preprocessing steps (e.g., global signal regression), and multilayer network analyses-related methodological decisions (e.g., window size, edge definition, number of optimizations) on reliability, as well as to optimize predictive accuracy.

## Acknowledgements

The work presented here was primarily supported by the NYS Office of Mental Health (AF, MPM, QT, SJC, ZY) and NIMH awards U01MH099059 (MPM), R24MH114806 (MPM), R01AG047596 (SJC, MPM), and R01MH111439 (MPM). Additional funding was provided by gifts to the Child Mind Institute (CMI) from Phyllis Green and Randolph Cowen. We would also like to thank the many individuals who have provided financial support to the CMI Healthy Brain Network to make the creation and sharing of this resource possible. SG was supported by the National Natural Science Foundation of China [NSFC-61876032]. CGY was supported by NSFC-81671774 and the Hundred Talents Program of the Chinese Academy of Sciences. Data were provided in part by the Human Connectome Project, WU-Minn Consortium (Principal Investigators: David Van Essen and Kamil Ugurbil; 1U54MH091657) funded by the 16 NIH Institutes and Centers that support the NIH Blueprint for Neuroscience Research; and by the McDonnell Center for Systems Neuroscience at Washington University.

## CRediT authorship contribution statement

**Zhen Yang**: Conceptualization, Software, Formal analysis, Visualization, Project administration, Writing - original draft; **Qawi K. Telesford**: Conceptualization, Software, Validation, Formal analysis, Writing - original draft; **Alexandre R. Franco**: Conceptualization, Resources, Writing - original draft; **Shi Gu**: Conceptualization, Software; **Ting Xu**: Conceptualization, Software; **Lei Ai**: Resources; **Ryan Lim**: Software, Data Curation; **Francisco X. Castellanos**: Conceptualization; **Chao-Gan Yan**: Conceptualization; **Stan Colcombe**: Conceptualization, Writing - original draft; **Michael P. Milham**: Conceptualization, Resources, Writing - original draft. All co-authors contributed to Writing-review & editing.

## Supplementary Materials

### Methods

#### Eriksen Flanker task

In this task, participants were asked to focus on the center arrow and indicate whether the arrow was pointing left or right by pushing a button with their left or right index finger. The task consisted of four types of trials with an event-related design: congruent (flanker arrows pointing in the same direction), incongruent (flanker arrows pointing in the opposite direction), neutral (flanker arrows replaced with diamonds), and no-go (flanker arrows replaced with X’s) trials. There was a total of 120 trials per type presented across three separate imaging runs. For each trial, the stimulus was presented for 1250 ms and the inter-trial intervals randomly varied among 1450, 2900, 4350, and 5800 ms.

#### Human Connectome Project

The test and retest data of 45 subjects who have retest imaging data were utilized (https://www.humanconnectome.org/study/hcp-young-adult/data-releases). Each subject had four resting state fMRI runs (∼15 min/run) collected using the same protocol on a 3T Siemens Skyra with a multiband gradient-echo EPI sequence at test and retest (multiband factor=8, TR=0.72 s, echo time = 33.1 ms, field of view = 208 × 180 mm^2^, number of slices = 72, voxel size = 2 mm^3^, and flip angle = 52°). All subjects are monozygotic twins (19 twin pairs and 7 without a paired twin). Three subjects were excluded due to incomplete data acquisition (total number of volumes less than 80%). For the remaining 42 subjects (16 twin pairs and 10 without a paired twin), one subject from each twin pair (the one who is more right-handed and/or with a test-retest interval closer to the median) was chosen. Only one subject from each twin pair was included to avoid the twin-related decrease in between-subject variance. Additionally, one unpaired twin who was left-handed was excluded, leaving a total number of 25 right-handed subjects for analysis (age: 22 to 35 years, 9 males/16 females, test-retest intervals ranging from 52 to 326 days).

Functional images were preprocessed identically as HBN-SSI data using CPAC. To explore the impact of preprocessing pipeline, the publicly released extensively processed data were also used. This enhanced pipeline included gradient distortion correction, motion correction, field bias correction, spatial registration into a common Montreal Neurological Institute (MNI) space, intensity normalization (Glasser et al. 2013), and artifact removal using independent component analysis FIX (Griffanti et al. 2014, Salimi-Khorshidi et al. 2014). Unlike the CPAC pipeline, global signal regression was not performed in the HCP pipeline. After preprocessing, we extracted mean signals from each ROI based on the CC200 atlas (Craddock et al. 2012) and applied a sliding window to time series (window length 139 TRs, ∼100 s what is comparable to which was used for the HBN-SSI dataset). For each window, edges were estimated using wavelet coherence. The dynamic community detection, the computation of dynamic network measures, and the estimation of test-retest reliability of dynamic network measures were performed in the same way as in the HBN-SSI data.

#### Simulated Data

To test the impact of the GenLouvain algorithm on recovering known underlying dynamic changes in community structure, we created a toy multilayer network dataset. It consisted of 128 nodes which were divided into four 32-node communities where each community represented a complete graph (i.e., all nodes were interconnected with each other). Across 13 layers, these four communities either split to form a 16-node community or merged to form a 32-node community. As shown in **Figure S3A**, the four communities split or merged at different rates; Community 1 did not change, Community 2 split or merged every three layers, Community 3 split or merged every two layers, and Community 4 split or merged every layer. The changes in community structure across layers (over time) were captured using the GenLouvain algorithm (**Figure S3B**). Whenever the community assignment of a node changed, this fact was recorded and used to calculate node flexibility; as shown in **Figure S3C**, node flexibility varied across nodes in the four original communities, with nodes that were originally in Community 1 showing no changes and nodes that were originally in Community 4 showing the most changes.

Given that output from the GenLouvain algorithm is nondeterministic, it is common practice to run the algorithm across multiple optimizations. In the case of the toy network, after 1000 optimizations, a change should occur with a 50% probability at each split or merge. This behavior occurs because all edges in the network have equal weight, and thus when a 32-node community splits, there is equal likelihood that the nodes forming the new community come from either set of 16 nodes. Likewise, when a community merges, there is equal likelihood that the older community will cease and join the new community, or that the newer community will cease and return to the older community.

## Figure legends

**Figure S1.**
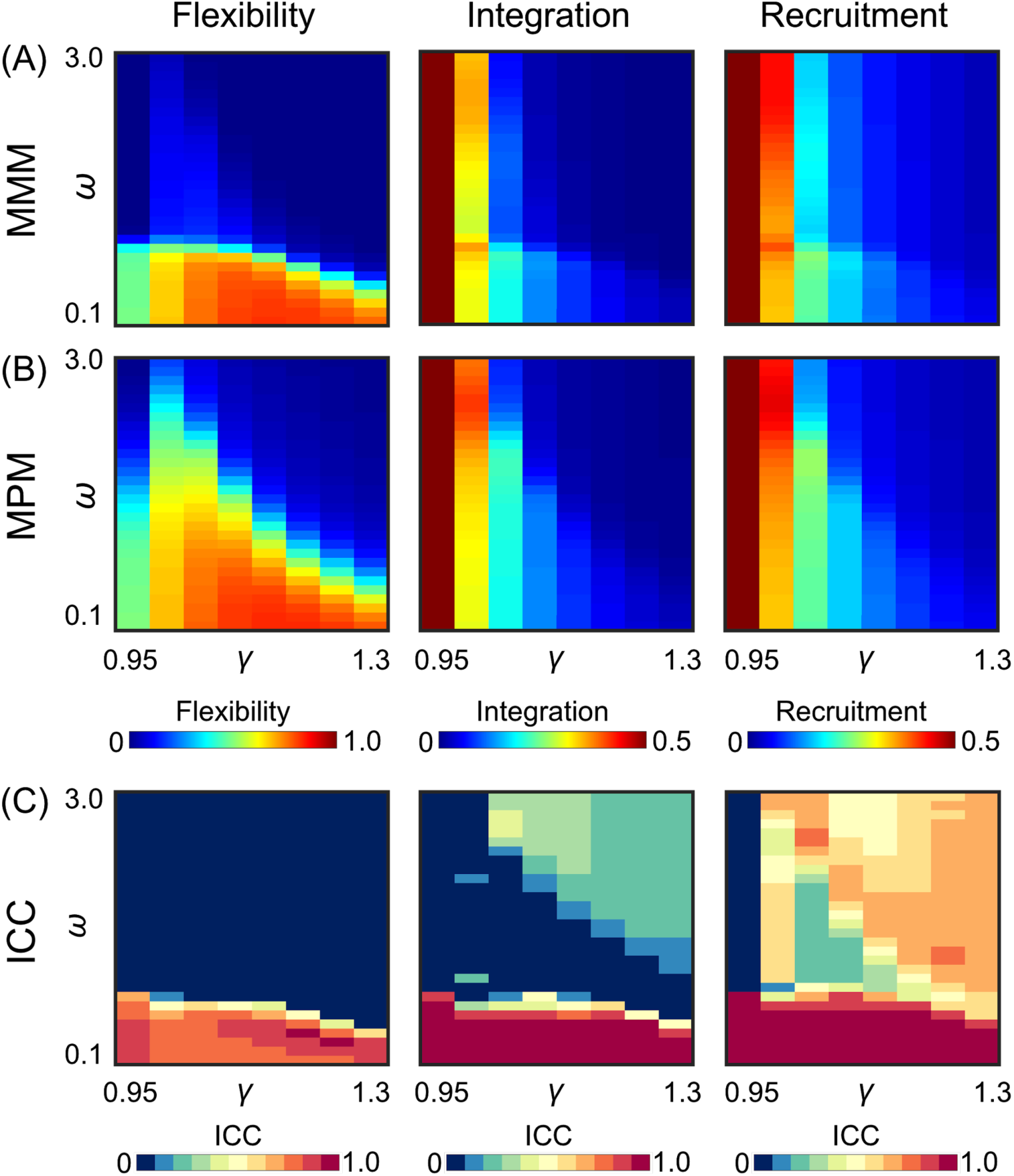
The impact of the generalized Louvain (GenLouvain) method on dynamic network measures. In the 2-dimensional γ−ω parameter space, abrupt discontinuity in estimated flexibility, integration, and recruitment values were observed when the Maximum Modularity Method (MMM) was used **(A)**. This issue was most serious for flexibility, and less so for integration and recruitment. The updated Modularity Probability Method (MPM) mitigated this issue and resulted in an apparently more continuous landscape **(B)**. Reliability between the two methods above the point of apparent discontinuity was close to zero for flexibility, ranged from poor to fair for integration, and ranged from poor to good for recruitment (**C**).

**Figure S2.**
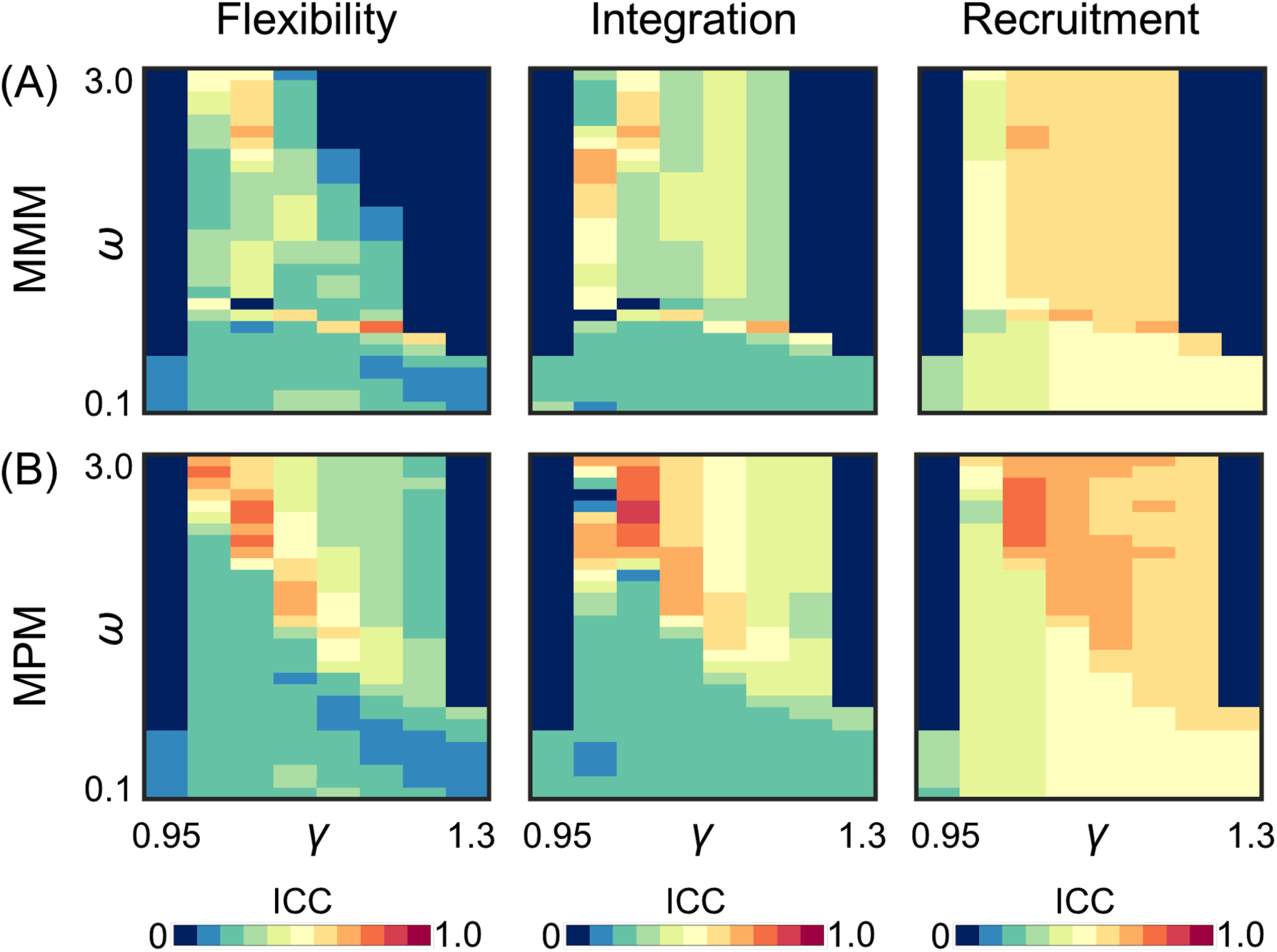
The test-retest reliability of dynamic network measures was impacted by the GenLouvain method. The test-retest reliability in the portion of the parameter space above the apparent discontinuity was lower for the default MMM (**A**) compared to the new MPM (**B**) for flexibility, integration, and recruitment. Results were obtained for the movie condition with 60 minutes of data.

**Figure S3.**
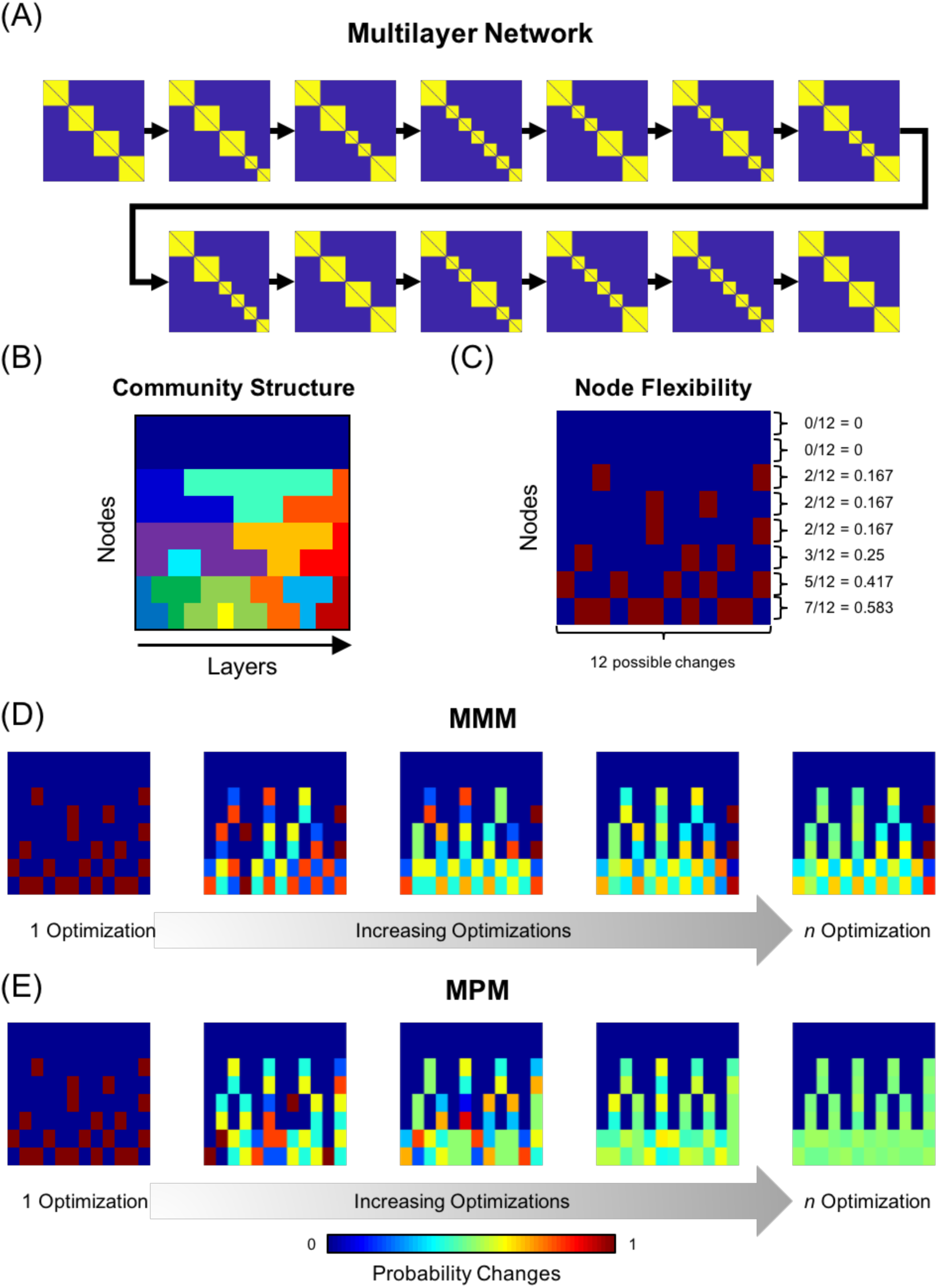
The impact of the GenLouvain method on multilayer network analyses in the toy data. (**A**) A multilayer network representing groups of nodes split into four communities showed the splitting and merging of communities across 13 layers. (**B**) Community structure across layers was identified by the GenLouvain algorithm (we showed one optimization here). (**C**) Node flexibility quantifies how often a node changes community assignment. From single optimization, flexibility was calculated by finding the number of times that a node changed a community divided by the number of possible times the nodes could change. In practice, flexibility is calculated across multiple optimizations. Using the GenLouvain algorithm across *n* = 1000 optimizations, it is expected that at the point where a community changes, there is a 50% chance that a group of nodes will form the new community or merge with an old community. When comparing method choice, it was readily apparent that although results appeared visually similar at one optimization, the MMM (**D**) did not result in community changes with equal likelihood at each split or merge, while the MPM (**E**) produced the expected outcome for this toy network.

**Figure S4.**
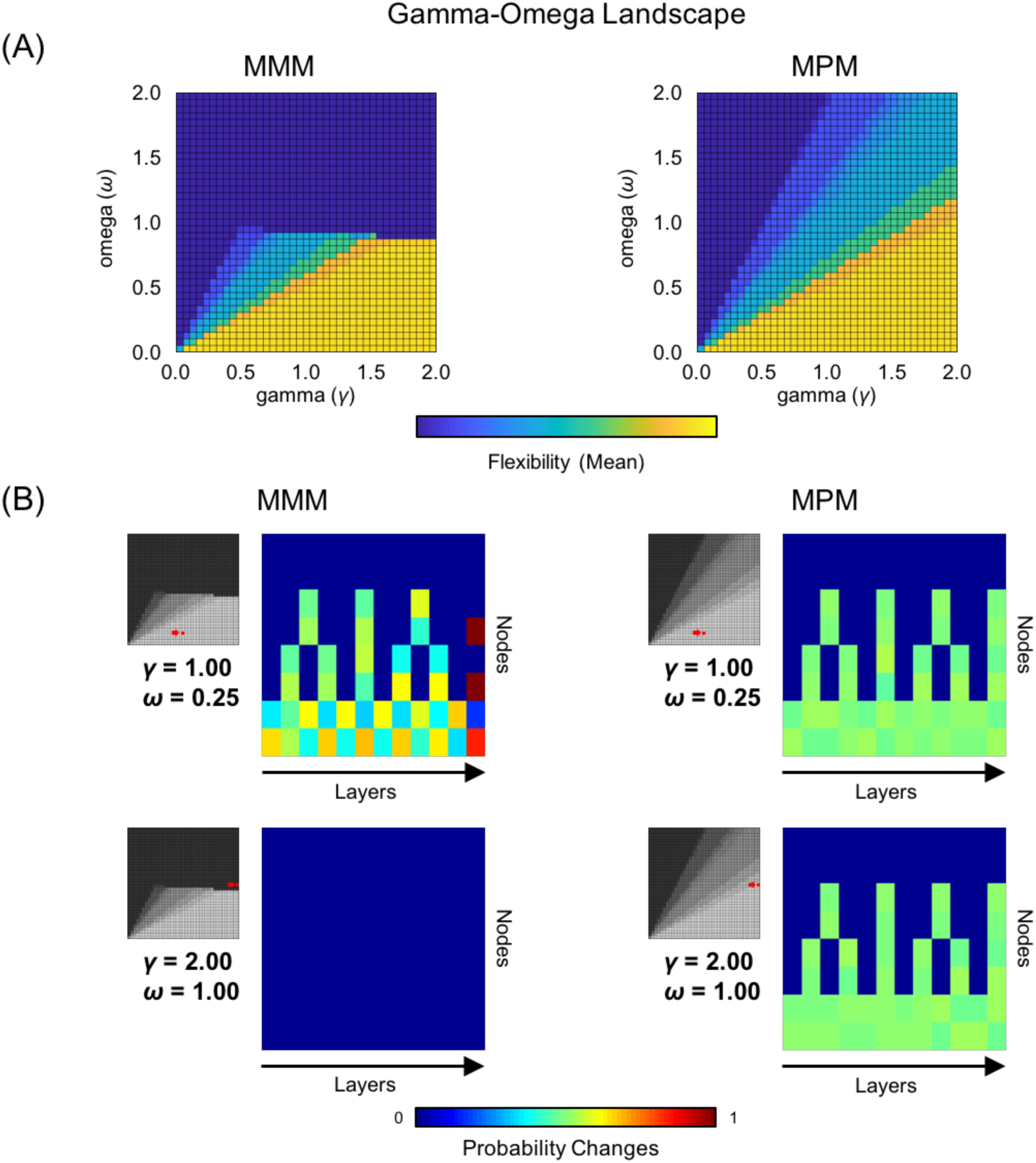
The MPM outperformed the MMM in terms of recovering the underlying dynamic changes in modular structure regardless of the γ−ω selection. When using the GenLouvain algorithm, the parameters for γ and ω changed the average flexibility measured across nodes. (**A**) MMM resulted in an abrupt drop off in flexibility values at ω > 1. In contrast, the newer method MPM resulted in an apparently more continuous modulation of the metric values. (**B**) Although the flexibility values in the parameter space appeared similar below the dropoff, multiple optimizations revealed stark differences in the converging results. When choosing a value below the apparent discontinuity (γ = 1.00, ω = 0.25), the output from the newer MPM method matched the expected outcome for the toy network. In contrast, the default MMM did not produce the expected outcome. When choosing a value above the apparent discontinuity, MPM was still able to recover the expected outcome while MMM did not find any changes.

**Figure S5.**
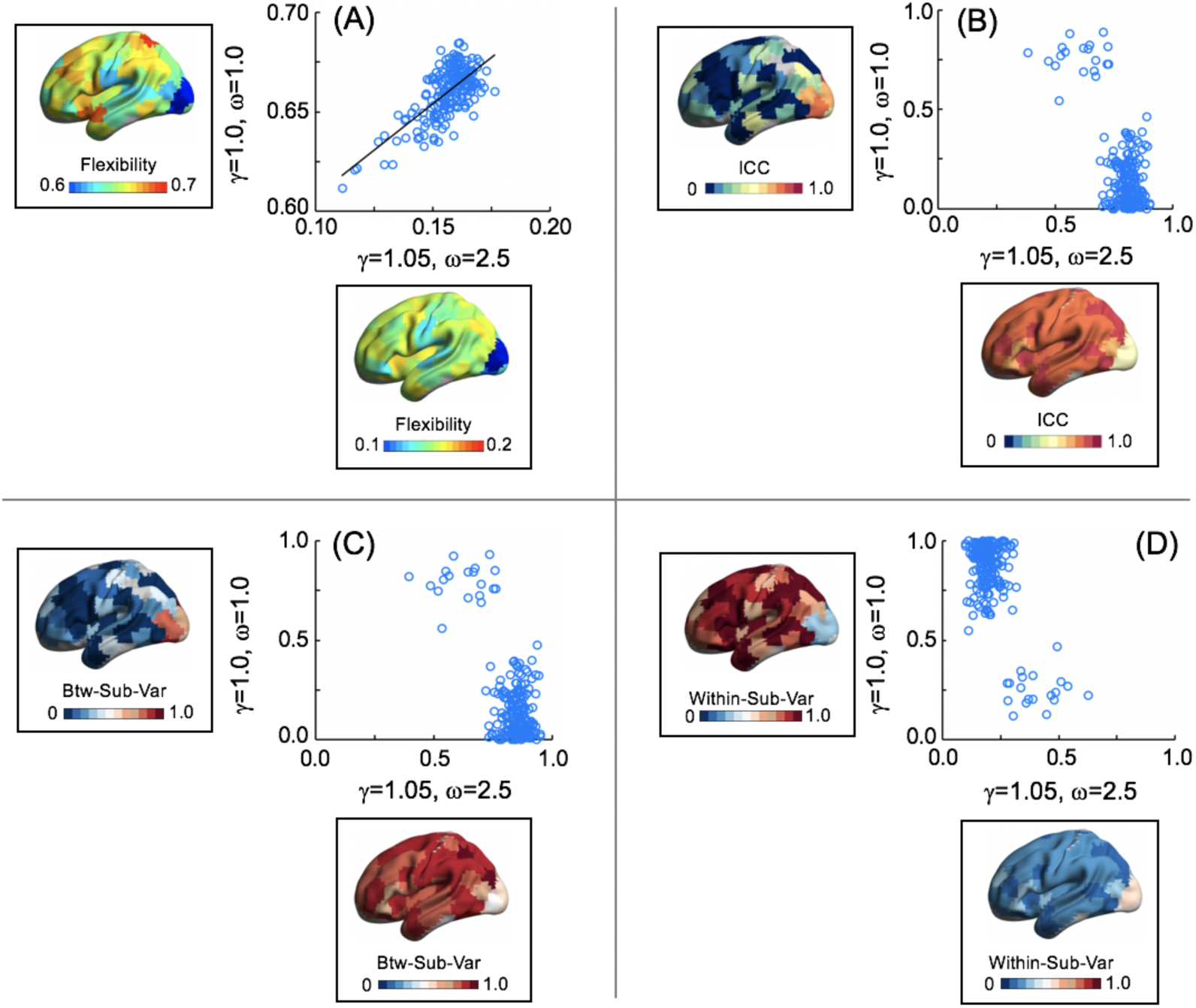
Comparison of our parameter selection [γ=1.05, ω=2.5] and previously recommended parameters [γ=1, ω=1]. The spatial topographies of mean flexibility computed across subjects for two parameter choices were similar (**A**: Pearson’s r=0.70), even though the range of flexibility values differed. Consistent with previous work (Betzel et al. 2017), we found that high-order cognitive regions had greater flexibility than primary cortices. For the movie condition, the visual cortex had the lowest flexibility. Compared to the parameter choice [γ=1, ω=1], [γ=1.05, ω=2.5] had a much higher test-retest reliability across the brain (except for the visual cortex) (**B**). The low test-retest reliability of the flexibility observed at [γ=1, ω=1] was driven by low between-subject variance (Between-Sub-Var: **C**) and high within-subject variance (Within-Sub Var: **D**). The relatively low ICCs (although still medium in size) observed in the visual cortex at [γ=1.05, ω=2.5] were associated with comparable within- and between-subject variance. In the scatter plot, each dot represents an ROI. These results were obtained based on the movie condition using 60 minutes of data.

**Figure S6.**
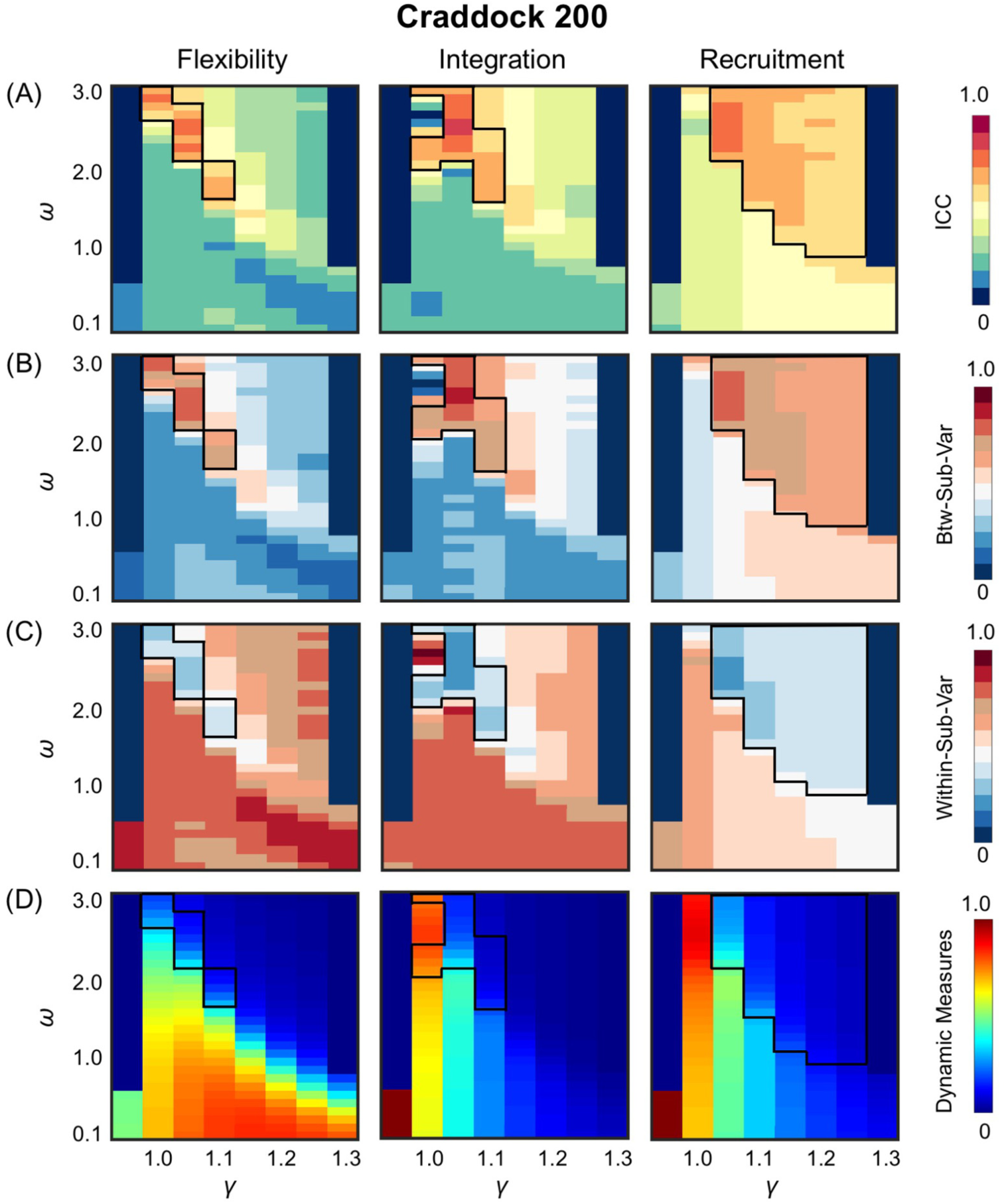
Characterization of the high ICC parameter space obtained in the main analyses using Craddock 200 parcellations. ICC values, between-subject variance (Btw-Sub-Var), within-subject variance (Within-Sub-Var), and global dynamic measures across the γ-ω plane were shown in **Panel A, B, C**, and **D** respectively. High ICCs (areas with ICC≥0.6 are indicated by black lines) overlapped largely with the portion with high Btw-Sub-Var and low Within-Sub-Var, as well as low global dynamic measures (integration has some exceptions). Results shown are the 60-min movie condition.

**Figure S7.**
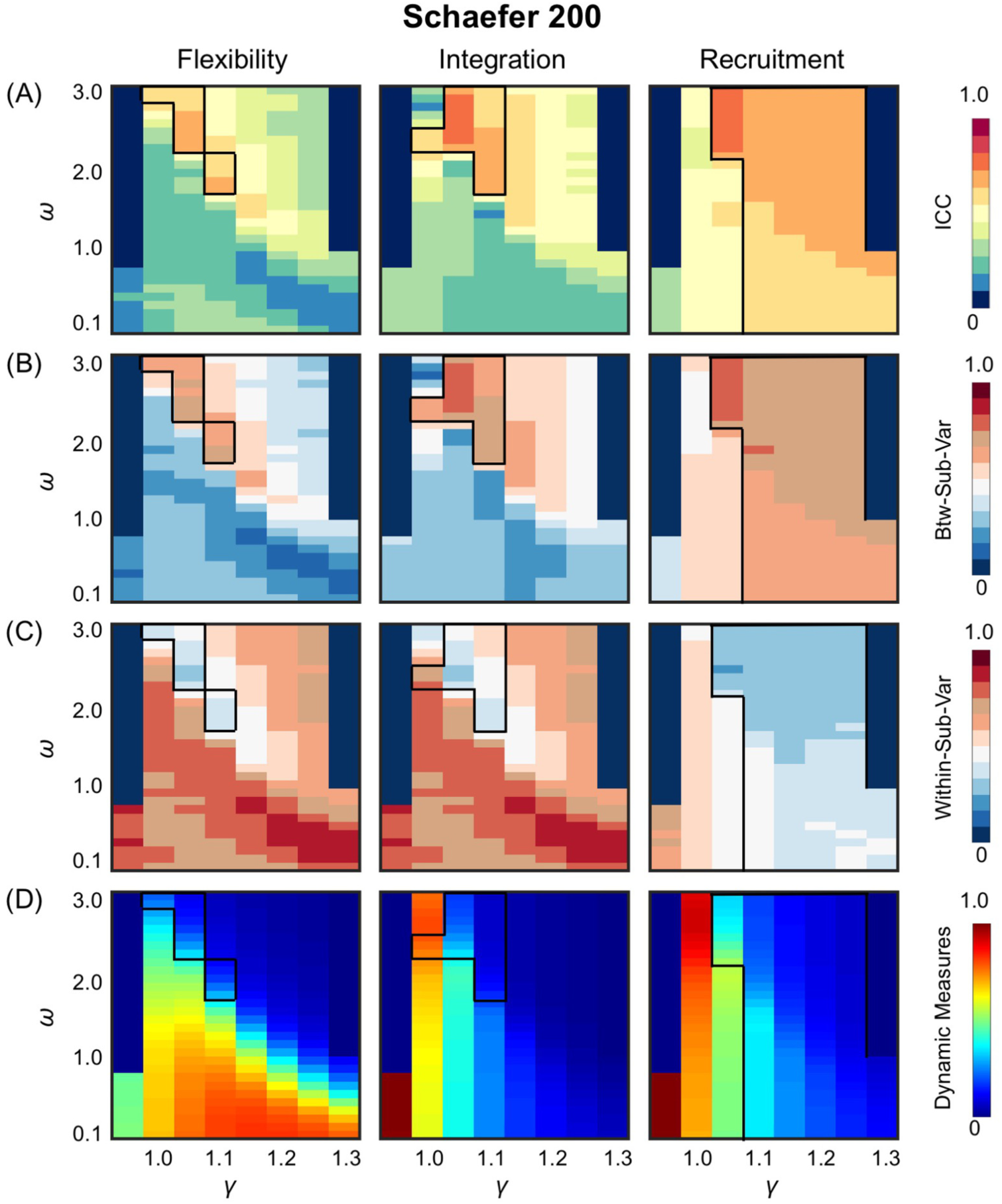
Characterization of the high ICC parameter space obtained using Schaefer 200. For each panel, the layout was the same as Figure S6.

**Figure S8.**
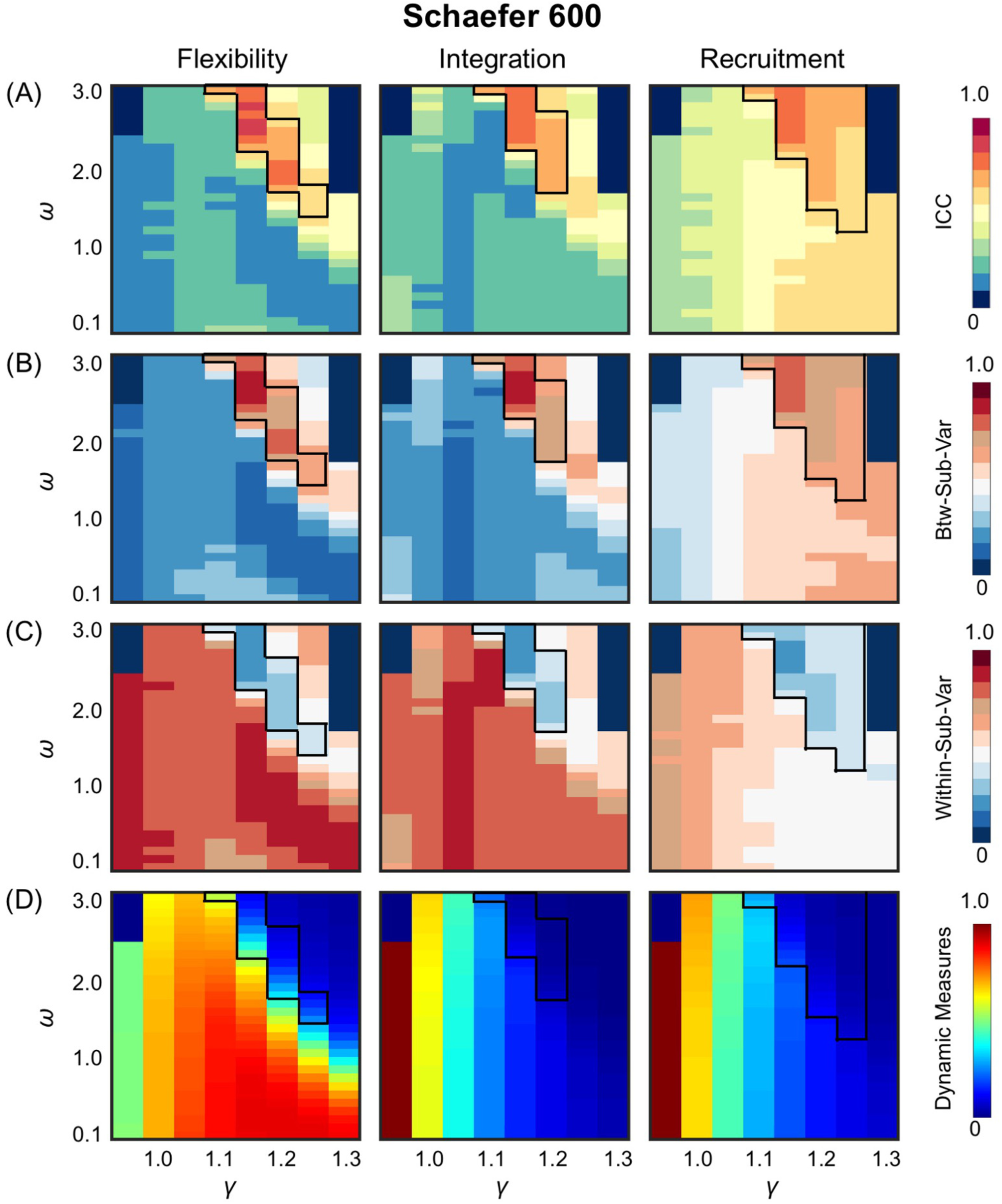
Characterization of the high ICC parameter space obtained using Schaefer 600. For each panel, the layout was the same as Figure S6.

**Figure S9.**
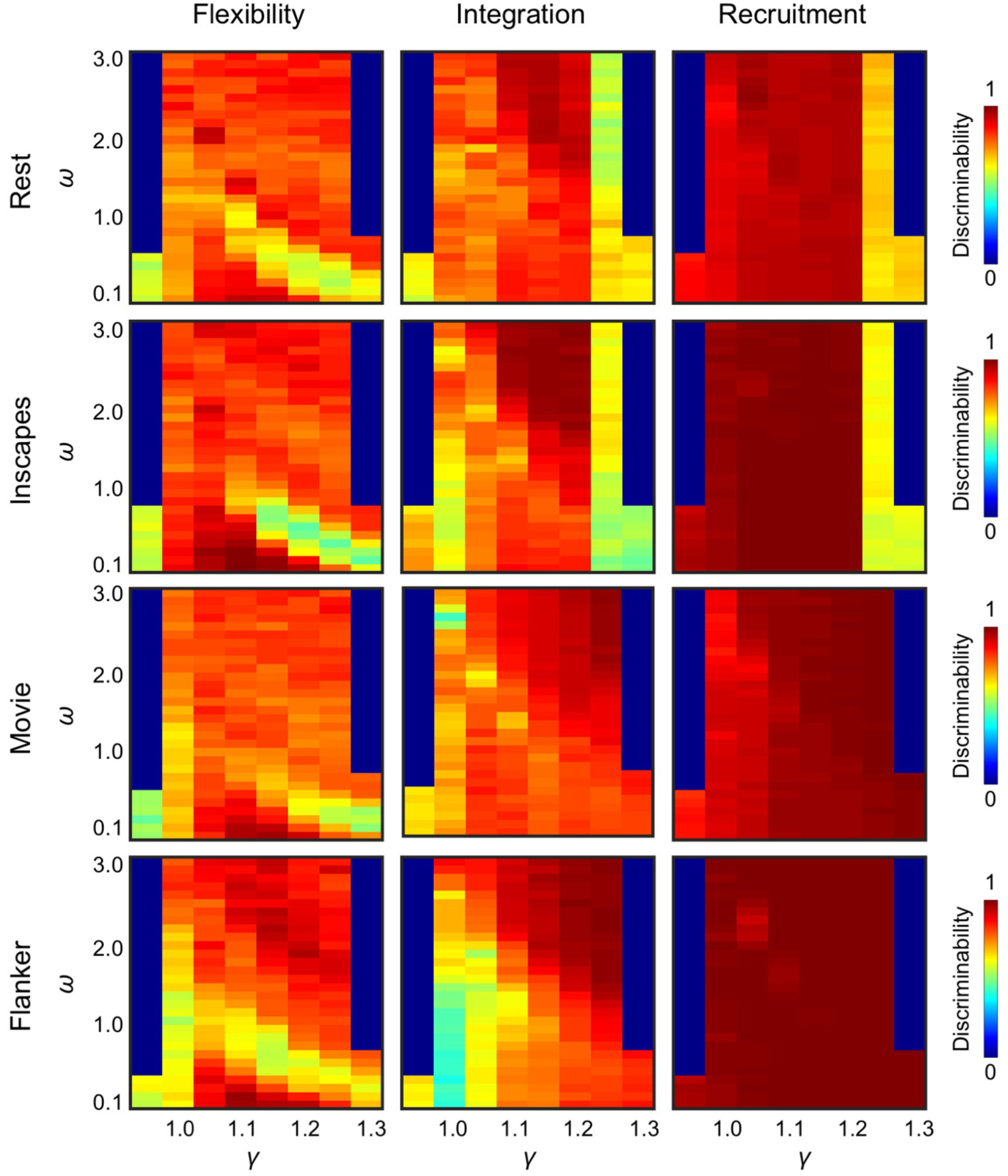
The discriminability of dynamic network measures depended on the γ-ω selection. In the γ-ω plane, global discriminability was computed across 200 ROIs. The range with high discriminability (>0.9) differed from the range with high test-retest reliability, especially for flexibility and integration. For recruitment, discriminability was high for most of the parameter space. For a given measure, global discriminability was less consistent across tasks (compare rows). Note that the values in the parameter space where the number of communities was smaller than 2 or greater than 100 were set to zero in each plane. Discriminability was evaluated with the maximal amount of data available (60 min).

**Figure S10.**
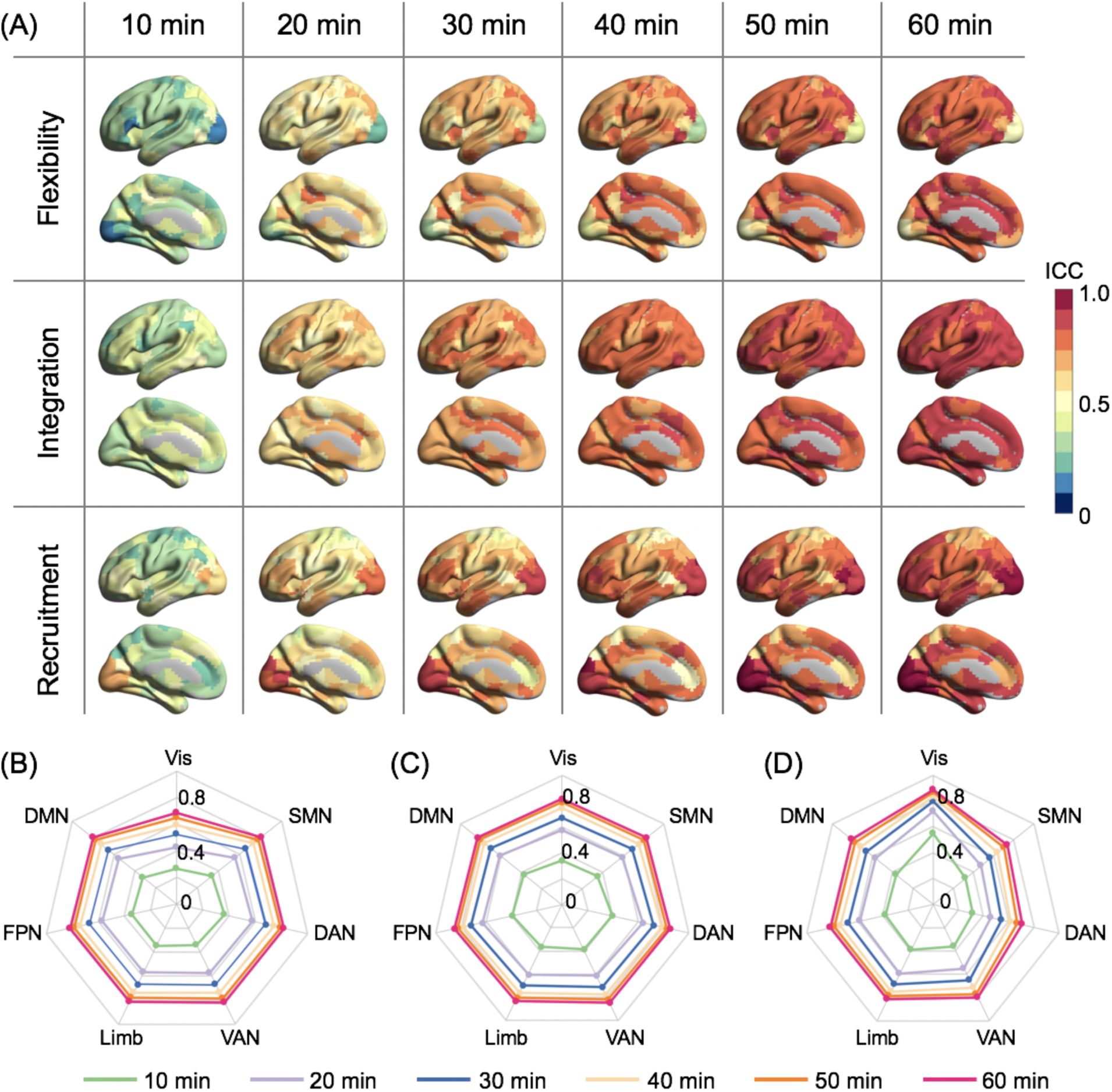
Regional and network-level variations in reliability improvement as a function of scan duration for the movie condition. (**A**) The ICC values were plotted on the brain’s surface for flexibility, integration, and recruitment at six scan durations. For ease of interpretation, regional ICC values were summarized using Yeo et al. (2011)’s seven networks for three measures (flexibility: **B**; integration: **C**; and recruitment: **D**) and each of the six scan durations. With 10 min of data, the ICC values of the dynamic metrics were low in all networks, except for recruitment in the visual network. With increased scan duration, the reliability of the dynamic metrics improved, and the improvement was most noticeable from 10 to 20 min. The visual network had the lowest reliability for flexibility and the highest reliability for recruitment. The reliability of recruitment within the somatomotor network was also relatively low. Spatial variation was less obvious for integration than for the other two measures.

**Figure S11.**
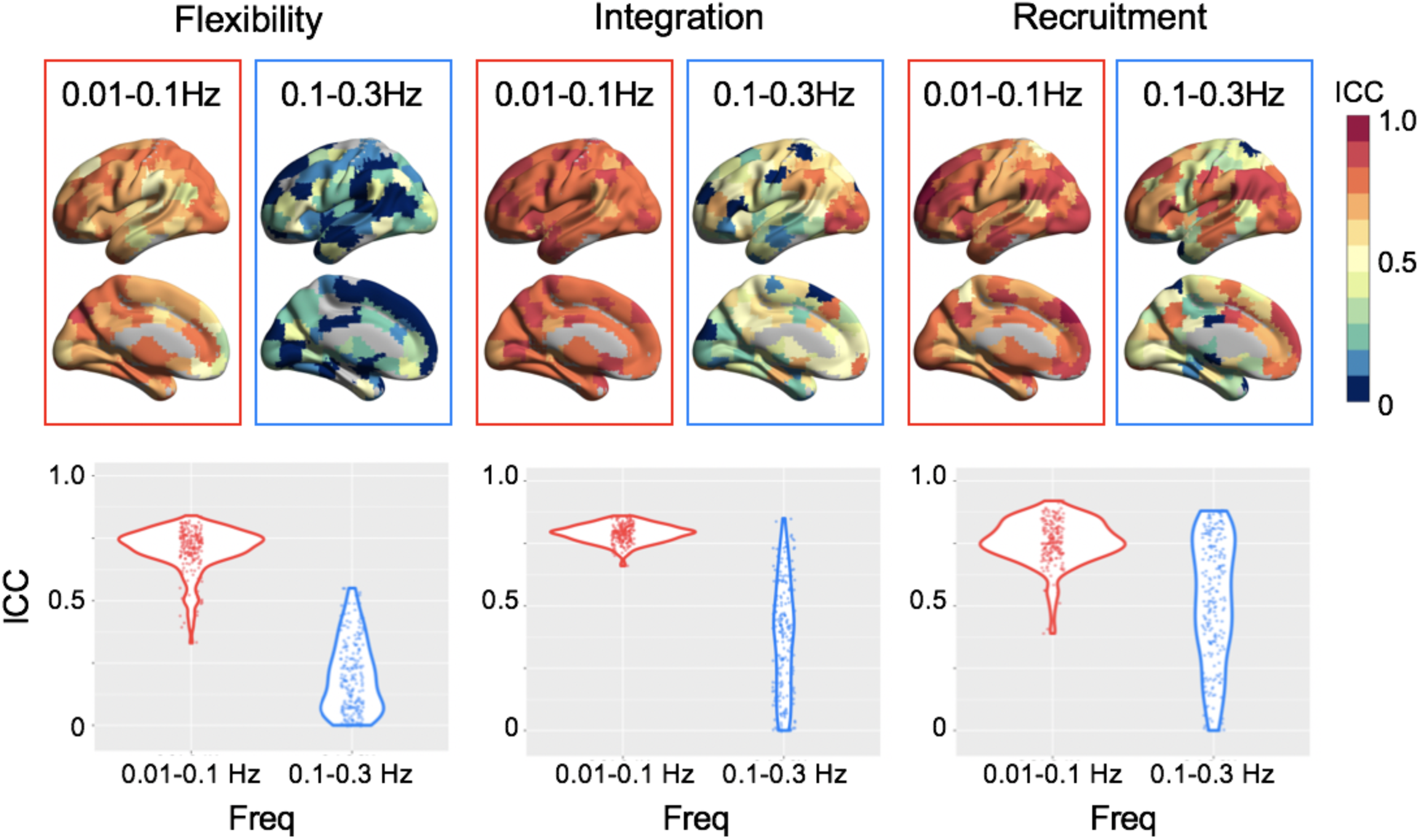
For the flanker task, dynamic network measures were more reliable when estimated from the low frequency components of the fMRI signal (0.01-0.1Hz) compared to the high frequency components (0.1-0.3Hz). ICCs of the 200 ROIs were plotted on the brain surface and summarized in a violin plot for low (red) and high (blue) frequency components of the fMRI signal.

## Notes

### Competing Interest Statement

The authors have declared no competing interest.

### Summary of Updates

In this updated version, we have made major revisions on results and conclusions.

